# Novel high-quality amoeba genomes reveal widespread codon usage mismatch between giant viruses and their hosts

**DOI:** 10.1101/2024.09.23.614596

**Authors:** Anouk Willemsen, Alejandro Manzano-Marín, Matthias Horn

## Abstract

The need for high-quality protist genomes has prevented in-depth computational and experimental studies of giant virus-host interactions. In addition, our current knowledge of host range is highly biased due to the few hosts used to isolate novel giant viruses. This study presents six high-quality amoeba genomes from known and potential giant virus hosts belonging to two distinct eukaryotic clades: *Amoebozoa* and *Discoba*. We employ their genomic data to investigate the predictability of giant virus host range. Using a combination of long and short-read sequencing, we obtained highly contiguous and complete genomes of *Acanthamoeba castellanii, Acanthamoeba griffini*, *Acanthamoeba terricola*, *Naegleria clarki*, *Vermamoeba vermiformis*, and *Willaertia magna*, contributing to the collection of sequences for the eukaryotic tree of life. We found that the six amoebae have distinct codon usage patterns and that, contrary to other virus groups, giant viruses often have different and even opposite codon usage with their known hosts. Conversely, giant viruses with matching codon usage are frequently not known to infect or replicate in these hosts. Interestingly, analyses of integrated viral sequences in the amoeba host genomes reveal potential novel virus-host associations. Matching of codon usage preferences is often used to predict virus-host pairs. However, with the broad-scale analyses performed in this study, we demonstrate that codon usage alone appears to be a poor predictor of giant virus host range. We discuss the potential strategies that giant viruses employ to ensure high viral fitness in non-matching hosts. Moreover, this study emphasises the need for more high-quality protist genomes. Finally, the amoeba genomes presented in this study set the stage for future experimental studies to better understand how giant viruses interact with different host species.

## Introduction

Protists are a diverse group of microorganisms representing various evolutionary lineages in the eukaryotic tree of life (Burki, 2014; Simpson et al., 2017). They are also defined as eukaryotes that are not animals, plants, or fungi, with most being single-celled organisms. Amoebae are polyphyletic (Burki et al., 2020) and are among the protist members that can migrate by a process known as amoeboid movement (Webb & Horwitz, 2003; Yoshida & Soldati, 2006). They are present in various environments, from soil and freshwater to marine habitats, and their extensive genetic diversity reflects their diverse ecological niches and lifestyles. Some amoebae are known to cause disease in humans. For example, certain members of the family *Acanthamoeba* are causal agents of a severe sight-threatening infection of the cornea (Lorenzo-Morales et al., 2015), and the so-called “brain-eating” amoeba *Naegleria fowleri* can cause a rare but nearly always fatal brain infection (Grace et al., 2015). Amoebae can be associated with various types of symbionts, such as bacteria (Horn & Wagner, 2004; Molmeret et al., 2005; Shi et al., 2021), algae (Weiner et al., 2022), viruses (Oliveira et al., 2019) and fungi (Corsaro et al., 2014; Steenbergen et al., 2001). Interaction with these symbionts can be mutualistic, parasitic, or commensal. With amoebae hosting such a variety of organisms, they constitute a favourable environment for genetic exchange between sympatric symbionts resulting in organisms with complex chimeric genomes (Boyer et al., 2009; Moliner et al., 2010; Moreira & Brochier-Armanet, 2008; Wang & Wu, 2017), such as giant viruses.

Giant viruses are a highly unusual group of viruses, of which the first member was discovered only two decades ago (La Scola et al., 2003; Raoult et al., 2004). These viruses are classified within the phylum *Nucleocytoviricota*, an assemblage of several families of double-stranded DNA (dsDNA) viruses infecting multicellular and unicellular eukaryotes. As their name suggests, giant viruses distinguish themselves from other viruses with vast particle and genome sizes (Abrahão et al., 2018; La Scola et al., 2003; Legendre et al., 2014; Philippe et al., 2013). The size of their virions exceeds that of the smallest known bacteria and archaea. Their large genomes revealed an unexpected complexity, including the existence of hundreds of genes that have not yet been attributed to viruses (Abrahão et al., 2018; Brahim Belhaouari et al., 2022; Needham, Yoshizawa, et al., 2019; Schulz et al., 2017). Interestingly, giant viruses do not form a monophyletic clade within the *Nucleocytoviricota* and appear to have evolved on multiple independent occasions from smaller viruses (Koonin & Yutin, 2018). While the evolutionary factors that promote genome expansion in giant viruses remain unclear, it is suggested that virus-host interactions may play an important role where viral size limitations imposed by multicellular hosts appear to restrict giant viruses to unicellular eukaryotic (*i.e.,* protist) hosts (Koonin et al., 2020). Yet, the natural host species and precise host range remain unknown for most giant viruses.

Our current knowledge of giant virus host range is strongly biased towards lytic viruses that have been isolated through co-cultivation with a limited number of protists, mainly with *Acanthamoeba* species (Abrahão et al., 2018; Boyer et al., 2009; La Scola et al., 2003; Legendre et al., 2014, 2015; Yoshikawa et al., 2019) but also with members of the genera *Vermamoeba* (Abrahão et al., 2018; Bajrai et al., 2016; Reteno et al., 2015), *Bodo* (Deeg et al., 2018), *Cafeteria* (Fischer et al., 2010), and more recently *Naegleria* (Arthofer et al., 2024). Yet, metagenomic studies suggest that *Acanthamoeba*-associated giant viruses are less prevalent under natural conditions. Rather, these studies suggest other amoeboid flagellates and algae species as natural hosts (Needham, Poirier, et al., 2019; Schulz et al., 2017; Zhang et al., 2015).

In several organisms, it has been shown that a correlation between codon usage and tRNA content exists (Anderson, 1969), where coevolution of codon usage and tRNA content can optimise the efficiency of translation (Higgs & Ran, 2008; Ikemura, 1985; Rocha, 2004). Most known viruses have highly compact genomes that do not encode tRNAs. Thus, the translation of viral proteins relies on the host tRNA pool. This situation creates translational selection for the adaptation of viral codon usage to those of their hosts, ensuring efficient viral translation. Therefore, the prediction of viable virus-host pairs is often based on similarities in codon usage. This method works well for phages, as there is generally a strong similarity in codon usage between prokaryotes and these viruses (Esposito et al., 2016; Lucks et al., 2008; Sau et al., 2005). However, for most eukaryotes (animals and plants), there appears to be an overall poor match of codon usage with their infecting viruses (Simón et al., 2021). Among protists, it was even noted that there is a negative correlation (Simón et al., 2021). For certain giant viruses, the presence of tRNAs and other translation-related genes in their genomes (Abrahão et al., 2018; Koonin & Yutin, 2018; Raoult et al., 2004; Schulz et al., 2017) might explain how these viruses can thrive well in protist hosts. Indeed, for one large DNA virus (*Ostreococcus tauri virus 5*) infecting a marine green alga (*Ostreococcus tauri*), it has been described that viral tRNAs complement the host tRNA pool for translational optimisation of the viral genes (Michely et al., 2013). Unfortunately, the lack of data does not allow for an accurate portrait of virus-protist interactions at the genomic level.

While high-quality genomic resources for diverse giant virus lineages are currently available, a significant gap exists regarding similar resources for their hosts. Thus, in this study, we sequence and provide high-quality genomes of six amoeba that we currently use in our laboratory to study their interactions with giant viruses: *Acanthamoeba terricola* Neff (up to recently classified as *Acanthamoeba castellanii* Neff (Corsaro et al., 2024)), *Acanthamoeba castellanii* 1BU, *Acanthamoeba griffini* Sawyer, *Vermamoeba vermiformis* CDC-19, *Naegleria clarki* RU30, and *Willaertia magna* T5(S)44. These amoebae belong to two distantly related eukaryotic clades: the *Amoebozoa* within the Amorphea supergroup and *Discoba* within the unresolved Excavates supergroup (Burki et al., 2020). Yet, despite their unrelatedness, most of these amoebae can host giant viruses, reflecting the remarkable structural and genomic diversity we observe among giant viruses (Fischer et al., 2023; Koonin & Yutin, 2018). To investigate whether codon usage plays a vital role in giant virus host adaptation, we compared the codon usage preferences of both partners. In addition, we also examine the amoeba genomes for viral integrations, as some of these have been shown to act as antiviral defence systems (Fischer & Hackl, 2016; Levasseur et al., 2016) and have the potential to reveal novel giant virus-host associations (Bellas et al., 2023; Maumus & Blanc, 2016). Finally, we evaluate additional important factors for predicting giant virus host range and discuss potential strategies that giant viruses employ to optimise viral fitness in their hosts.

## Results and Discussion

### The six amoeba genomes are highly contiguous and complete

We used a combination of Illumina short reads and Oxford Nanopore long reads to assemble the amoeba genomes (see Materials and Methods). For all six amoebae, the genome assemblies were contained in a low number of scaffolds (Table 1), with 90% of the individual genomes contained within 57 (*A. terricola* Neff), 123 (*A. castellanii* 1BU), 280 (*A. griffini* Sawyer), 132 (*V. vermiformi*s CDC-19), 559 (*W. magna* T5(S)44) and 670 (*N. clarki* RU30) scaffolds. These are the first published genome assemblies of *A. castellanii* 1BU, *A. griffini*, *N. clarki*, and *W. magna* T5(S)44. For *V. vermiformis* and *W. magna,* the assemblies significantly improve the number of contigs and contiguity compared to previously published assemblies of the same species (Table S1). Compared to the *A. terricola* Neff reference assembly, this assembly is slightly longer with fewer contigs (Table S1). Recently, a somewhat longer (43.8 Mb) *A. terricola Neff* assembly was published, where with the inclusion of Hi-C, the assembly resulted in 111 scaffolds (Matthey-Doret et al., 2022).

**Table 1.** Genome assembly statistics for the final assemblies of the six protist genomes.

| Species | Size (Mbp) | Scaffolds (No.) | N50 (Mbp) | L50 (No.) | L90 (No.) | Largest sequence (Mbp) | Total GC% | protein-coding genes (No.) |
| --- | --- | --- | --- | --- | --- | --- | --- | --- |
| *Acanthamoeba terricola* Neff | 42.97 | 246 | 0.9 | 17 | 57 | 1.9 | 58.42 | 14924 |
| *Acanthamoeba castellanii* 1BU | 47.37 | 474 | 0.5 | 25 | 123 | 2.0 | 58.53 | 14761 |
| *Acanthamoeba griffini* Sawyer | 52.91 | 862 | 0.4 | 33 | 280 | 1.5 | 56.66 | 16497 |
| *Vermamoeba vermiformis* CDC-19 | 43.35 | 309 | 0.4 | 39 | 132 | 0.8 | 41.94 | 15385 |
| *Naegleria clarki* RU30 | 53.71 | 1709 | 0.1 | 109 | 670 | 1.2 | 32.27 | 22696 |
| *Willaertia magna* T5(S)44 | 47.61 | 1239 | 0.09 | 140 | 559 | 0.6 | 24.69 | 18644 |

Genome completeness was estimated based on single-copy orthologs using BUSCO scores (Simão et al., 2015). These were calculated for the genome assemblies and the genome annotations at the protein and transcript levels. The completeness score for the genomes varied between 78.1% and 86.6% of complete eukaryotic universal single-copy orthologs (Table S2). For most genomes, these scores improved notably when considering the annotated proteins (Table S2) and transcripts (Fig. 1 and Table S2), with for the transcripts scores of 94.1% (*V. vermiformi*s), 91.4% (*A. terricola* Neff, *A. griffini*), 91.0% (*A. castellanii* 1BU), 84.3% (*W. magna*) and 81.6% (*N. clarki)*. Only the annotated *W. magna* and *N. clarki* genomes appear to be missing some BUSCO gene matches that are, in fact, present in the genome (Table S2), as their genome assembly completeness scores are higher (86.6% and 85.1%, respectively). This incongruency reveals the limitations of the gene prediction tool we used for amoebae in the *Discoba* clade.

**Figure 1.**
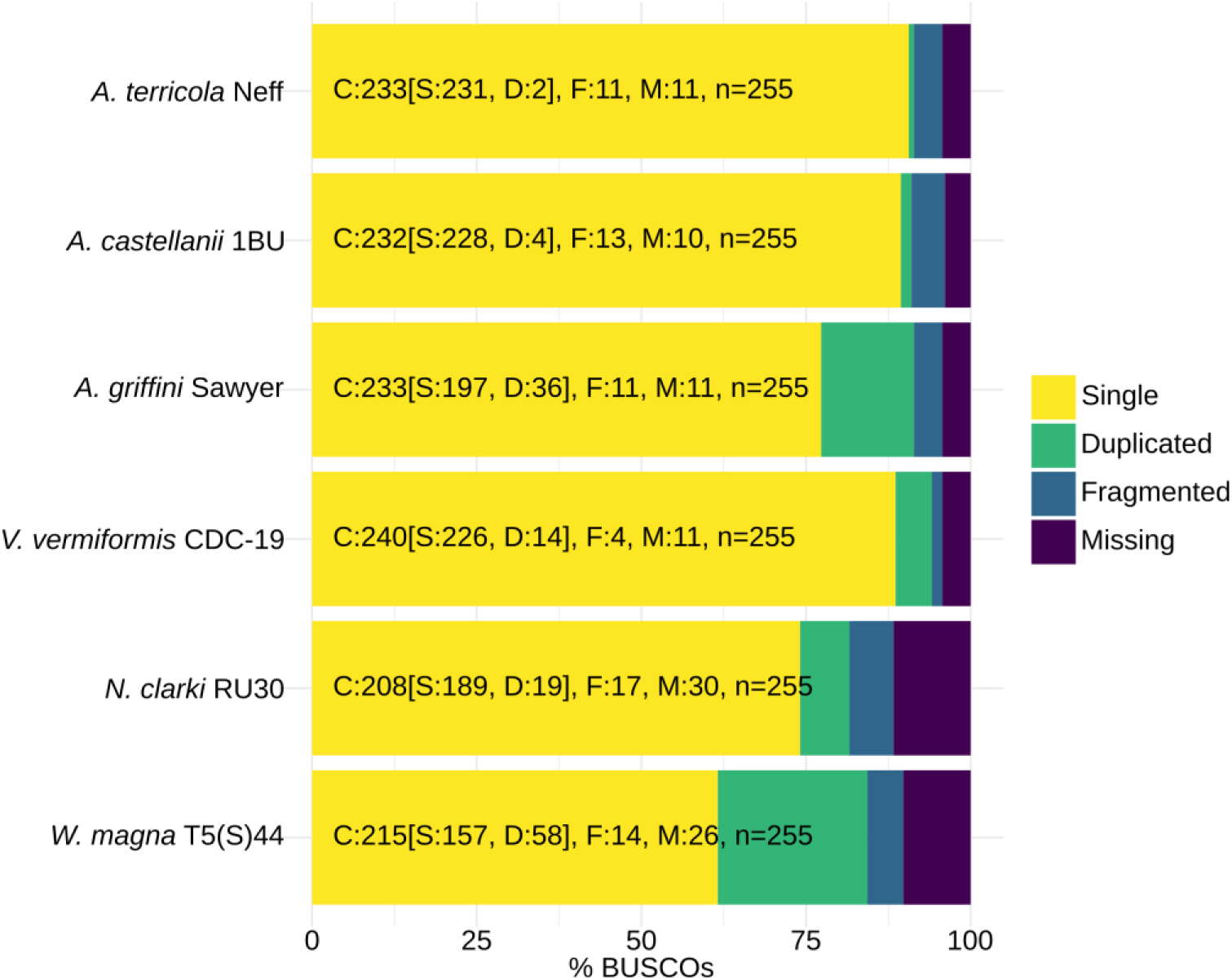
Genome completeness (BUSCO scores) for the annotated genomes of the six amoebae. BUSCO was run using the transcriptome mode against the eukaryotic database (C: complete [D: duplicated], F: fragmented, M: missing, n: total BUSCO groups searched). These results indicate high completeness levels of between 81.6% and 94.1%.

### Phylogenetic and phylogenomic analyses define amoeba relationships

From each genome sequenced in this study, 18S rRNA sequences were extracted and aligned with other complete and near-complete protist 18S rRNA sequences. Phylogenetic trees were constructed for the phyla *Discosea* (Fig. S1; including *Acanthamoeba* spp.), *Tubulinea* (Fig. S2; including *V. vermiformis*) and *Heterolobosea* (Fig. S3; including *N. clarki* and *W. magna*). These trees demonstrate that on the 18S rRNA level, the amoebae cluster within their expected clades with their close relatives. Nonetheless, 18S rRNA gene sequences are insufficient to resolve amoeba at the species level as the identity between closely related strains is high (Supplementary Data at Zenodo repository). Despite the high 18S rRNA sequence identity, a previous study has already shown considerable variation in the mitochondrial protein-coding genes and was thus able to uncover diversity even between conspecific *Acanthamoeba* and *Vermamoeba* (Fučíková & Lahr, 2016). We identified a unique region in the mitochondrial genome of *W. magna* T5(S)44 sequenced in this study, containing two hypothetical proteins shared with *Naegleria* spp., allowing us to distinguish this strain from other *W. magna* strains (Supplementary Data at Zenodo repository). These results stress the need for more high-quality complete genome sequences to perform genome-scale analyses and resolve relationships among amoebae.

To contribute to the collection of sequences for the eukaryotic tree of life, we used the package PhyloFisher (Tice et al., 2021) and its associated database for phylogenomic analyses. The resulting phylogenomic trees of the *Amoebozoa* (Fig. 2A) and *Discoba* (Fig. 2B) eukaryotic clades show high levels of bootstrap support, where the *Acanthamoeba*, *Vermamoeba*, and *Naegleria* sequences form separate monophyletic clades, and *Willaertia* appears as a sister taxon to the two available *Naegleria* species. The two *A. terricola* Neff genomes that derive from the same isolate are not considered identical in the phylogenomic tree (Fig. 2A). This is not unexpected as the reference genome of this isolate (RefSeq: GCF_000313135.1) has a higher representation of orthologous protein sequences (193 out of 228; Table S3), as compared to the *A. terricola* Neff genome sequenced in this study (188 out of 228; Table 3) in the manually curated database for tree construction (Materials and Methods). Nonetheless, since the genome sequenced in this study is retained in a lower number of scaffolds, future manual curation of the genome annotation will improve its ortholog representation for tree construction. The other amoeba genomes from this study have a much higher representation of orthologs in the tree construction database (between 198 and 224 out of 228; Table 3), comparable to other genomes in the database (Table S3), which, together with the high bootstrap values suggests strong support for their position in the phylogenomic tree (Fig. 2).

**Figure 2.**
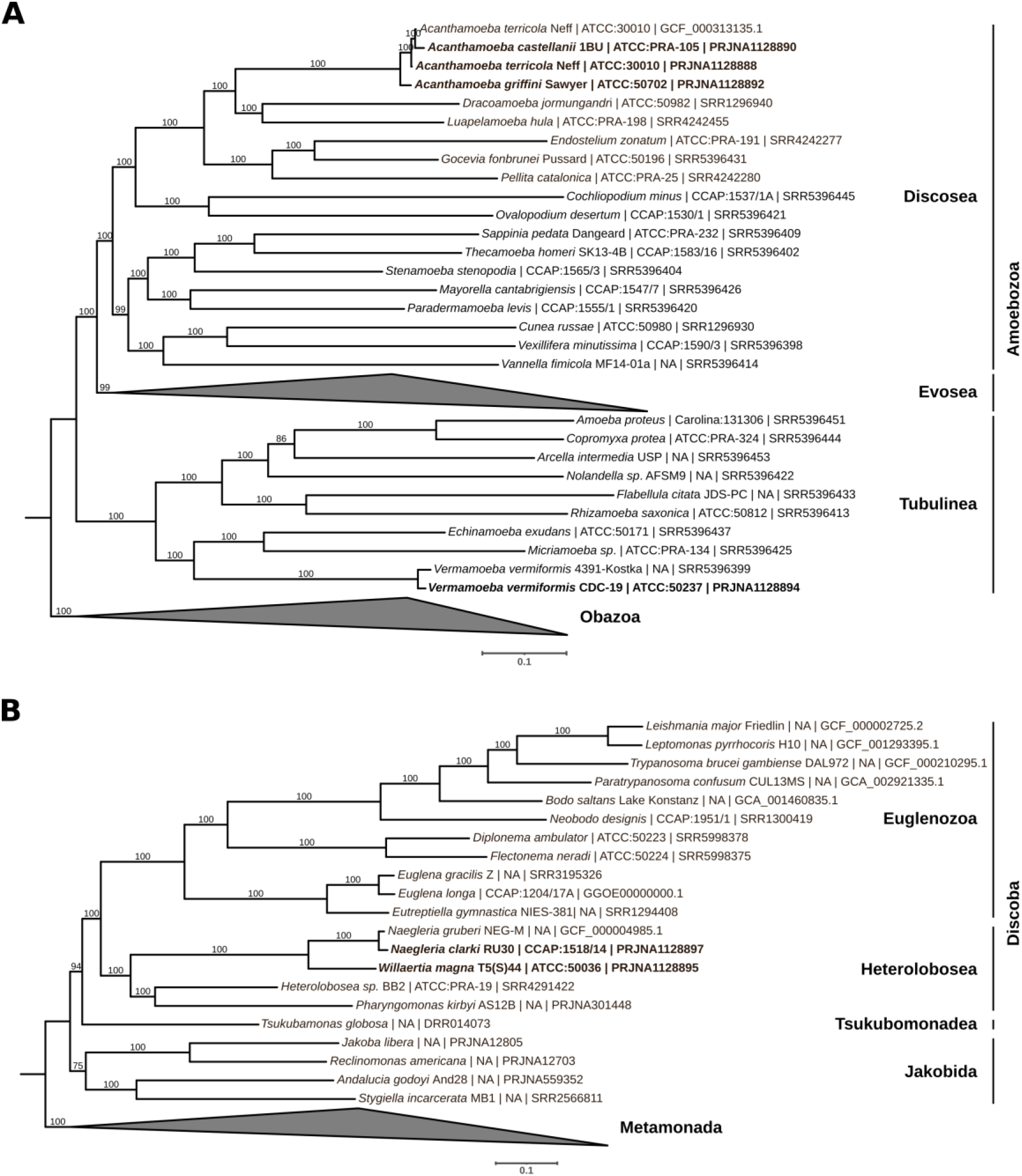
Phylogenomic trees inferred for two eukaryotic clades that include the newly sequenced amoebae. (**A**) This maximum likelihood phylogenomic tree for *Amoebozoa* was inferred from 228 proteins and 52 taxa, using the LG+R7 model of evolution. The names of the clades representing different phyla within the *Amoebozoa* are indicated on the right side. Members representing different phyla within the *Obazoa* clade (6 taxa) were used as an outgroup. (**B**) This maximum likelihood phylogenomic tree for *Discoba* was inferred from 228 proteins and 24 taxa, using the LG+F+R5 model of evolution. The names of the clades representing different phyla within the *Discoba* are indicated on the right side. Members representing different phyla within the *Metamonada* clade (3 taxa) were used as an outgroup. For both trees, bootstrap support values are given on the branches, and the names on the leaves are composed of “species names [strain] | culture collection database:accession number | NCBI dataset accession number” (see Table S3). The newly added taxa are indicated in bold font.

### Distinct codon usage patterns among the six amoebae

Before the generation of the high-quality genomes in this study, we were only able to perform codon usage analysis of *Acanthamoeba terricola* Neff (RefSeq: GCF_000313135.1), *Dictyostelium discoideum* AX4 (RefSeq: GCF_000004695.1) and three *Naegleria* species (RefSeq: GCF_000004985.1; GCF_008403515.1; GCF_003324165.1) that are not *N. clarki* (Fig. S4). Although *Dictyostelium* is a potential (Abrahão et al., 2018) and *Naegleria* a recently discovered novel giant virus host (Arthofer et al., 2024), we are unquestionably lacking high-quality genomes of other amoebae as currently most publicly available genomes in are simply too fragmented to generate reliable genome assemblies and annotations.

Using the additional five amoeba genomes generated here, we applied different methods to compare their codon usage. First, we analysed the relationship between the effective number of codons used in a gene (ENC) and G + C content in the third codon position (GC3). We used these values to construct ENC plots, allowing for intraspecific and interspecific comparisons of codon usage patterns (Wright, 1990). The ENC plots in Figure 3 show that the six amoebae have distinct codon usage patterns at the genus level. The three *Acanthamoeba* strains have a similar GC3 composition, with most codons being G- or C-ending. Opposite to *Acanthamoeba*, *N. clarki* and *W. magna* have low GC3 values. *V. vermiformis* has a considerable variation in GC3 values, ranging from 0.23 to 80, most likely reflecting variation in mutational bias among different regions of the genome. Except for *V. vermiformis*, the other five amoebae have a clear GC3 preference, while all six amoebae have a wide range of ENC values. Genes with low ENC values are often highly expressed (Mohasses et al., 2020; Wright, 1990)(blue data points in Figure 3), as these genes usually use the minimal subset of codons that are recognised by the most abundant tRNA species (Puigbò et al., 2007). These results suggest that for the six amoebae, besides mutation, there is translational selection acting for the usage of preferred codons by highly expressed genes.

**Figure 3.**
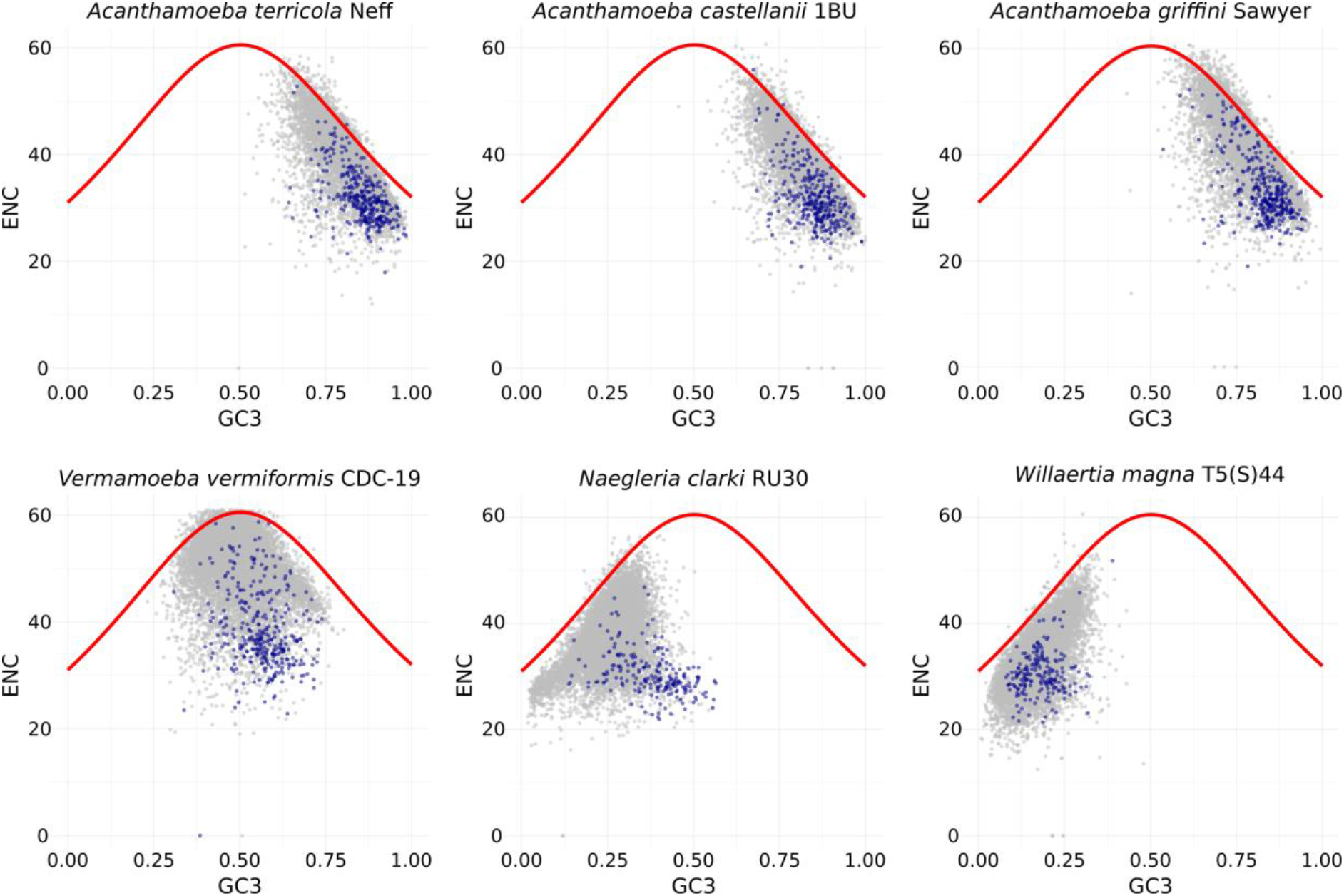
Distinct codon usage patterns across six amoebae genomes. ENC values were plotted against GC content at the third codon position (GC3). Each gray or blue dot represents a gene (*A. terricola*: *N*=11217, *A. castellanii*: *N*= 11903, *A. griffini*: *N*=13504, *V. vermiformis*: *N*=14170, *N. clarki*: *N*=21306, *W.magna*: *N*=18192). The blue dots represent the top 5% of fragments per kilobase million (FPKM) values from the corresponding transcriptome assemblies as a proxy for identifying highly expressed genes. The continuous red curves represent the relationships between ENC and GC3 under the null hypothesis of no translational selection. If a particular gene lies on the red curve, it is suggested that it is subjected to mutational bias only (*i.e.* G+C compositional constraints). These plots show that the six protists have distinct codon usage patterns at the genus level (*i.e. Acanthamoeba, Vermamoeba, Naegleria* and *Willaertia*).

### Distinct codon usage preferences between giant viruses and their known hosts

We then investigated whether codon usage preferences could be used to computationally predict the host range of giant viruses. To be able to compare the codon usage preferences for viruses to those of the six amoebae, we calculated the Codon Adaptation Index (CAI) (Puigbò et al., 2008; Sharp & Li, 1987) and COdon Usage Similarity Index (COUSIN) scores (Bourret et al., 2019). Using these two indices, we compared all available full-length giant virus genomes (Tables S4) at both the family (Fig. 4) and genus (Fig. S5-S10) levels to each host. We generally observed a high correlation between the CAI and COUSIN scores (Table S5). Since the COUSIN scores allow for comparison between organisms (Bourret et al., 2019), we only show the COUSIN_59_ scores in the main figures.

**Figure 4.**
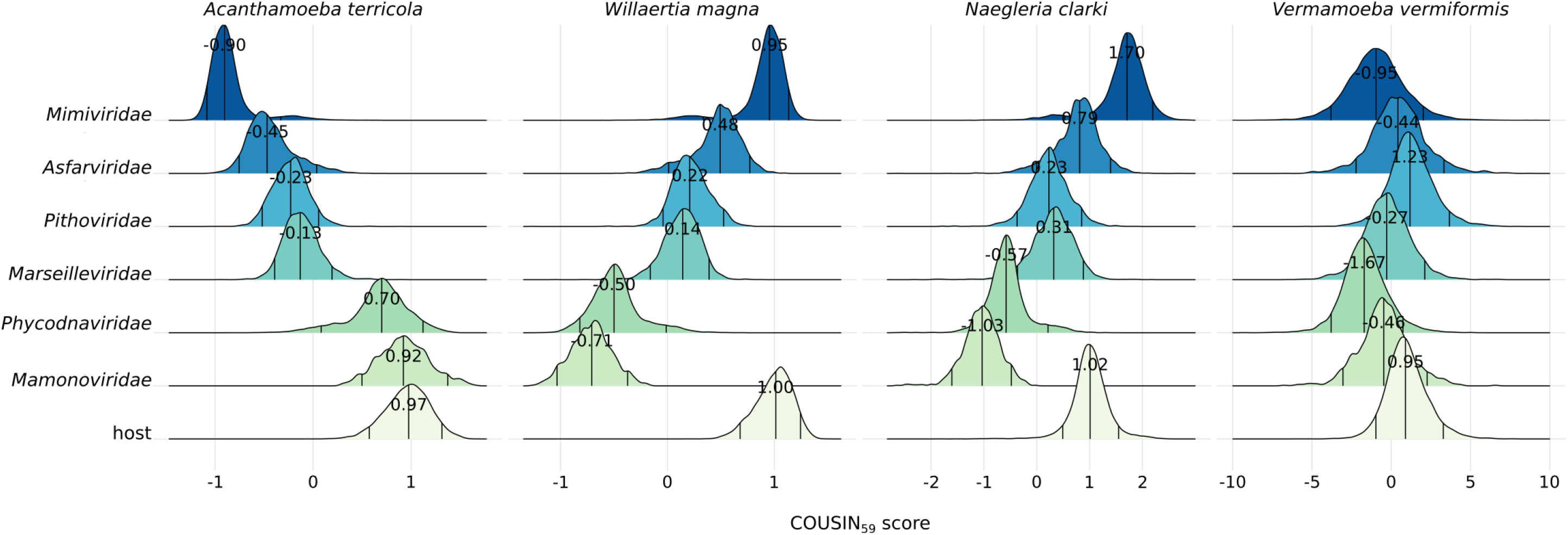
Density curves of the COUSIN_59_ score for giant viruses relative to their known and possible hosts. The density curves are organised by viral family (*Mimiviridae*: *N_viruses_*=29, *N_CDS_*=28969, *Asfarviridae*: *N_viruses_*=9, *N_CDS_*=4292, *Pithoviridae*: *N_viruses_*=6, *N_CDS_*=3393, *Marseilleviridae*: *N_viruses_*=10, *N_CDS_*=5735, *Phycodnaviridae*: *N_viruses_*=11, *N_CDS_*=9236, *Mamonoviridae N_viruses_*=2, *N_CDS_*=890). The bottom density curve always indicates the scores for the host, the name of which is indicated on top of each plot. The three lines within the density curves indicate the 95% confidence interval. The numbers within each curve indicate the centre values estimated by the Huber M-estimator of location. The COUSIN scores can be interpreted as follows: a score of 1 indicates that the codon usage preferences of viruses are similar to those of the corresponding host; a score of 0 indicates that there is equal usage of synonymous codons; above 1 indicates that codon usage preferences are similar but of larger magnitude (meaning that the codons that are most frequently used in the host are used even more frequently in the virus); between 0 and 1 indicates codon usage preferences are similar but of smaller magnitude (meaning that the codons that are less frequently used in the host are used even less frequently in the virus); below 0 means that the codon usage preferences of viruses are opposite to those of the corresponding host. Since the results for all three *Acanthamoeba* species were similar, only the results for *A. terricola* Neff are shown here. Note that the COUSIN_59_ score for *N. clarki* and *V. vermiformis* is depicted on a different scale for visibility.

The most intriguing result of this analysis is that viruses belonging to the *Mimiviridae* family (Huber M-estimator of location: −0.898, Median Absolute Deviation or MAD: 0.123) have opposite codon usage preferences to their best-known host *A. terricola* Neff (Huber-M: 0.968, MAD: 0.224; location difference = 1.862, *p* < 2.2e-16; Table S6; Fig. 4), whereas *Mimiviridae* (Huber-M: 0.947, MAD: 0.126) have close (but still significantly different) codon usage preferences to *W. magna* (Huber-M: 1.001, MAD: 0.170; location difference = 0.064, *p* < 2.2e-16; Table S6; Fig. 4), a potential (Abrahão et al., 2018) but yet unknown giant virus host. When we follow the theory that there is translational selection for adaptation of viral codon usage to those of their hosts, our results suggest that Mimiviruses have low fitness in *Acanthamoeba* hosts and high fitness in *W. magna*. However, Mimiviruses have been mainly isolated with *Acanthamoeba* hosts, and attempts to isolate giant viruses with *W. magna* have been unsuccessful so far (Boudjemaa et al., 2020). As reported previously, the *W. magna* genome contains genes related to viral sequences, with the majority being sequences from members of the *Mimivirdae* family (Hasni et al., 2019). This points towards horizontal gene transfer (HGT) events and past infections of mimiviruses in *W. magna*. However, the only mimivirus known to establish a productive infection in *W. magna* is *Tupanvirus soda lake* (Abrahão et al., 2018). Nevertheless, the viral titer increase over a 24-hour time period is six times lower in this host than in *A. terricola*. *Tupanvirus soda lake* has the largest translational apparatus (including a full set of tRNAs) within the known virosphere (Abrahão et al., 2018), suggesting that Tupanvirus presumably depends less on the host translation system as compared to other viruses. Therefore, it is unsurprising that Tupanvirus can thrive well in *A. terricola*. However, for Tupanvirus and other mimiviruses (with less encoded tRNAs), it remains to be investigated how exactly the translation-related genes are involved in coping with codon usage differences with their host(s).

Another intriguing result is that members of the *Mimiviridae* family (Huber-M: 1.697, MAD: 0.294) have codon usage preferences that are similar but of larger magnitude (*i.e.* codons that are most frequently used in the host are used even more frequently in the virus) to *N. clarki* (Huber-M: 1.015, MAD: 0.264; location difference = −0.684, *p* < 2.2e-16; Table S6; Fig. 4, Fig. S9) and other *Naegleria* host species (Fig. S4). This indicates that mimiviruses are super-optimised to *Naegleria,* where theory suggests efficient viral replication and high gene expression levels in this host. A recent study showed that a novel giant virus isolate (*Catovirus naegleriensis*, family: *Mimiviridae*), isolated with *N. clarki* as a bait, is specific to *Naegleria* host species, and does not induce infection phenotypes in the “typical” Mimivirus hosts *A. terricola* and *V. vermiformis* (Arthofer et al., 2024). Interestingly, *Catovirus naegleriensis* is only able to infect *Naegleria* host species under xenic conditions (*i.e.* with bacteria as a food source) but not under commonly used axenic conditions (Arthofer et al., 2024). For all future giant virus isolation studies, more natural culture conditions should be considered, as they can drastically change the infection outcome and have the great potential to reveal novel and natural giant virus hosts.

### Integrated viral sequences in the host genomes reveal potential novel virus-host associations

It has been recently shown that large parts of protist genomes are of viral origin and that most of these viral integrations appear to be functional viruses (Bellas et al., 2023). These endogenous viral elements (EVEs) comprise virophages, Polinton-like viruses and related entities (Bellas et al., 2023; Bellas & Sommaruga, 2021) and are comparable to prophage integrations in bacterial genomes. Although EVEs in eukaryotic genomes were previously thought to be self-synthesizing transposons (Kapitonov & Jurka, 2006), the detection of virus hallmark genes (*e.g.* capsid proteins and packaging ATPases) now suggest that many of these are endogenous viruses (Barreat & Katzourakis, 2021; Krupovic et al., 2014; Starrett et al., 2021).

In a previous study, endogenous virus MCP sequences were found in the assemblies of different *Acanthamoeba* species (*A. healyi*, *A. lenticulata*, *A. lugdunensis*, *A. mauritaniensis*, *A. pearcei*, *A. polyphaga*, *A. quina*, *A. rhysodes*, *A. royreba*), including two *A. castellanii* strains (Namur and astronyx), but not in the two *A. terricola* Neff strains that were interrogated (WGS accession: AHJI01000000 and AEYA01000000) (Bellas et al., 2023). The assemblies of *V. vermiformis* isolate TW EDP1, different *Naegleria* species (*N. fowleri*, *N. gruberi*, *N. lovaniensis*) and *W. magna* were also examined, but also here, no endogenous MCP sequences were found. Yet, most of these genomes have been generated with short-read data only, making the detection of EVEs challenging as they are often hidden in repetitive and difficult-to-assemble regions.

The long read data produced in this study facilitated the detection of MCP sequences in *Acanthamoeba*, as we found five MCP sequences integrated in *A. terricola* Neff, two in *A. castellanii* 1BU and three in *A. griffini* using DIAMOND BLASTX (Buchfink et al., 2015) and profile Hidden Markov Model (HMM) based searches (Tables S7 and S8). We did not detect any integrated MCP sequences in the *V. vermiformis*, *W. magna* and *N. clarki* genomes, which might well reflect a bias towards the few giant viruses represented in our database that are known to infect these hosts. Of all detected EVEs (Tables S7 and S8), four demonstrated a notable difference in GC content at the site of insertion compared to the genomic GC content (Figure 5). These differences are only apparent if integrated viruses have different codon usage preferences compared to those of their hosts. The inserted viral regions were flanked by terminal inverted repeats, supporting the hypothesis that these are genuine viral insertions. Interestingly, all four EVEs shown in Figure 5 gave a hit against yet unnamed viral MCPs identified through a previous analysis of all protist assemblies in the Genbank Whole Genome Shotgun database (Bellas et al., 2023). We confirmed these genes as MCPs by modelling their protein structures using AlphaFold. All four MCPs gave the best hit against the virus major capsid protein of *Paramecium bursaria Chlorella virus* type 1 (PBCV-1) (Nandhagopal et al., 2002). This virus infects the green algae *Chlorella* that can reside within the protist *Paramecium bursaria*, and has not (yet) been found associated with *Acanthamoeba* spp. The positive AlphaFold hits against PBCV-1 may simply reflect that this is one of the few *Nucleocytoviricota* members for which a high-resolution MCP structural reference is available (Fang et al., 2019; Shao et al., 2022). Other detected EVEs gave reliable BLAST and HMM hits against the MCPs of *Acanthamoeba castellanii medusavirus* and *Mollivirus kamchatka* (Tables S7 and S8). Medusaviruses (*Mamonoviridae*) and molliviruses (*Phycodnaviridae*) have close codon usage preferences to *Acanthamoeba* spp. (Fig. 4, Fig. S5-S7, Table S6), and are known to infect *A. terricola* Neff. However, our analyses suggest that HGT events have also occurred between these viruses and other *Acanthamoeba* spp., suggesting these as possible alternative common hosts.

**Figure 5.**
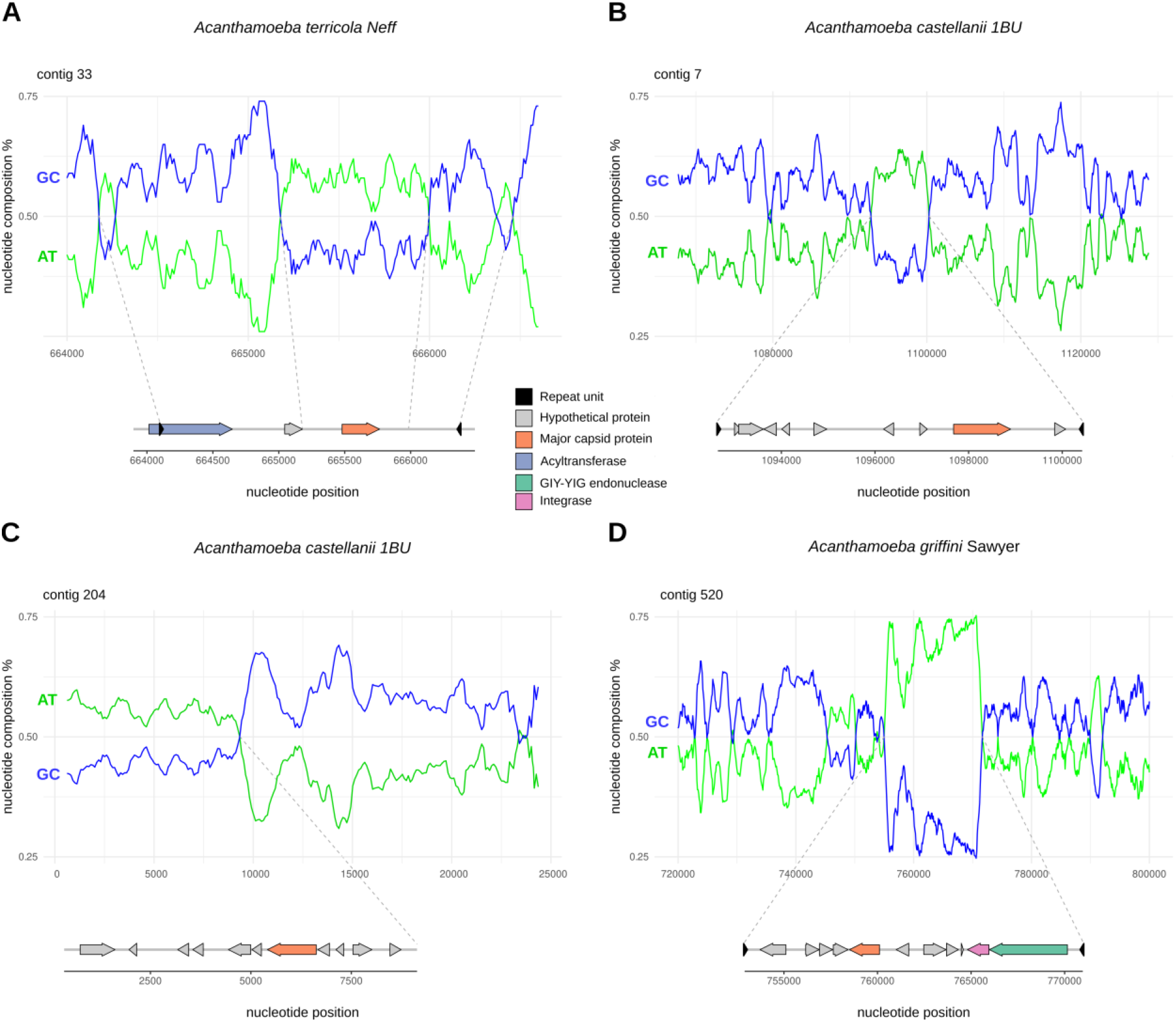
Examples of integrated viral sequences in the *Acanthamoeba* spp. genomes. In these cases, the integrated viral regions exhibit a notable difference in GC content from the host genome (A, B, C, D). They are flanked by terminal inverted repeats (A, B, D). The integrated viral regions shown here all contain viral major capsid proteins of unknown origin. The graphs show nucleotide position (bp) in each contig on the x-axis and GC (blue) and AT (green) percentages on the y-axis. Below each graph, a gene plot and predictions for the integrated viral region are shown. The contig in panel C is incomplete; therefore, only a part of the integrated viral region was detected, and thus, the detection of terminal inverted repeats was not possible.

Not all MCPs detected in this study appear to be intact genes, where stop codons and frameshifts suggest pseudogenisation of these. Only 2/5 integrated MCP genes in *A. terricola* Neff and 2/3 in *A. griffini* seem intact, whereas in *A. castellanii* 1BU this number is 0/2. While in most cases, error correction of long-read assemblies with short-read data significantly improves the consensus sequence, it is possible that in endogenous viral regions, this method causes genes to look artificially fragmented (Bellas et al., 2023). We, therefore, checked our Nanopore long-read assemblies before error correction with the Illumina short-read data. The observed stop codons and frameshifts in the detected MCP genes were still present, suggesting that these are no artefacts and that some of the integrated MCP genes have been degraded (Table S7). From the examples shown in Figure 5, the only intact MCP is the one detected in the integrated viral region in *A. griffini* (Fig. 5D). The presence of a retroviral integrase and a GIY-YIG endonuclease in the same region, plus the detection of intact and partial homologous MCP copies in other amoeba genomes (Table S7), suggests that this viral region can still actively move within and between genomes.

## Conclusion

The little information we currently have about the natural host range of giant viruses and the lack of high-quality host genomes is a hurdle for studying virus-host interactions. This study presents six amoeba genomes, of which five are known giant virus hosts. By comparing codon usage preferences of viruses and hosts, we demonstrate that this measure alone is not a good indicator for predicting giant virus host range. While it has already been reported previously that certain mimiviruses have highly dissimilar codon usage preferences to those of their host *A. castellanii* (Colson et al., 2013), this is the first study that performs a broad-scale analysis including all giant viruses with available full-length genomes, at the time of analyses, and their potential amoebal hosts, for which the high-quality genomes sequenced in this study where required. Indeed, we also find that the currently best-studied giant virus family (*Mimiviridae*) has codon usage preferences opposite those of their best-known laboratory hosts (*Acanthamoeba* spp.). However, this mismatch is not restricted to mimiviruses: our analyses reveal a widespread codon usage mismatch between giant viruses and their hosts. Despite this mismatch, giant viruses can maintain high viral fitness in these hosts. While for most giant viruses, alternative good matching hosts remain to be elucidated, the opposite also seems to occur; a good match in codon usage preferences can also result in low viral fitness (*e.g*., Tupanvirus in *W. magna*). Therefore, different giant viruses must have adopted different strategies to replicate and maintain high viral fitness in their mismatching hosts.

Notably, the extent of codon usage adaptation of viruses cannot be solely explained by a simple adaptation to the codon usage of their hosts as it reflects a combination of multiple selective and mutational pressures. For example, the host immune system also plays an important role, where immune defences drive viral codon usage away from sequences detected by the host (Lin et al., 2020). The replication site, nuclear or cytoplasmic, is also an important determinant of viral codon usage. Nuclear viruses (such as medusa-, molli- and pandoraviruses) tend to have a higher GC content for efficient nuclear export (Mordstein et al., 2020). However, most of the giant viruses we know to date replicate in viral factories within the cytoplasm of their host cells (such as mimi-, marseille- and pithoviruses) and do not have this selective pressure for a higher GC content. Therefore, these viruses may be able to afford such a conflict in codon usage with their hosts.

Interestingly, many viruses – including giant viruses – induce translational shutdown of their hosts (Abrahão et al., 2018; Bercovich-Kinori et al., 2016; Hsu et al., 2021). For phages, it has been shown that this phenomenon influences the selection on viral codon usage, leading to changes in demand for specific tRNAs during the course of infection and driving the acquisition of these tRNAs in viral genomes (Yang et al., 2021). In addition, the presence of translation-related genes may be a good strategy for giant viruses to sub-optimize their codon usage preferences and thereby avoid the host immune response. However, the translation-related genes present in viral genomes are not necessarily directly involved in compensating for codon usage differences with their hosts. Another strategy described for phages is to use tRNAs as a viral defence system, where phage-encoded tRNAs counteract tRNA-depleting strategies employed by enzymes from the host to defend from viral infection (van den Berg et al., 2023). The presence of few to many tRNAs in certain giant virus genomes (Koonin & Yutin, 2018) could help these viruses escape mutational pressures to adapt their codon usage preferences to that of their hosts.

While predicting virus-host pairs for giant viruses remains challenging, we can get good indications of viable pairs when taking into account additional information. Apart from codon usage preferences, the presence of viral integrations into the host genomes and host integrations into the viral genomes are good indicators of at least past interactions. The amoeba genomes presented in this study set the stage for future experimental studies to understand better how giant viruses interact with their hosts, bringing us a step closer to understanding the natural host range of giant viruses and their determining factors.

## Materials and Methods

### Strains and growth conditions

Amoebae were cultured in 75 cm^2^ and 175 cm^2^ culture flasks (Thermo Scientific cat nos. 156472 and 159920). *Acanthamoeba terricola* Neff (ATCC 30010) and *Acanthamoeba castellanii* 1BU (ATCC PRA-105) cells were grown axenically in Peptone-Yeast Extract-Glucose medium (PYG: ATCC Medium 712) at 25°C. *Acanthamoeba griffini* Sawyer (ATCC 50702) cells were grown axenically in PYG medium at 28°C. *Vermamoeba vermiformis* CDC-19 (ATCC 50237) cells were grown axenically in Serum Casein Glucose Yeast extract medium (SCGYEM: ATCC medium 1021) at 25°C. *Willaertia magna* T5(S)44 (ATCC 50036) cells were grown axenically in PYG medium plus 10% Fetal Bovine Serum (FBS: Gibco cat. no. 26140079) at 30°C. *Naegleria clarki* RU30 (CCAP 1518/14) cells were grown monoxenically with *Escherichia coli* (strain JW5503-1 *ΔtolC732::kan*) in Page’s Amoeba Saline buffer (PAS: ATCC medium 1323) at 25°C. *E. coli* was cultured in lysogeny broth medium (LB: 10 g/l NaCl, 10 g/l tryptone, 5 g/l yeast extract) at 37°C overnight with shaking at 200 rpm. *E. coli* was stored at 4°C upon usage.

### Nucleic acid extractions and genome sequencing

For DNA extraction, the amoeba cells were collected by centrifugation for 5 minutes at 10000✕g. The cells were washed with PAS. High-molecular-weight (HMW) DNA was extracted using the Wizard® HMW DNA Extraction Kit (Promega cat. no. A2920) according to the protocol for plant tissue with the following specifications/modifications: in step 3 the incubation at 65℃ was done for 25 minutes; in step 4 5µL of RNAse A was added, after mixing incubation at 37℃ was done for 25 minutes; in step 5 40µL of Proteinase K Solution was added, after mixing incubation at 56℃ was done for 25 minutes. For each amoeba, 3 to 5 individual HMW DNA extractions were combined. The DNA was further purified using Agencourt® AMPure® XP beads (Beckman Coulter cat. no. A63882) for subsequent long-read sequencing with the Oxford Nanopore Sequencing Technology. A test run was first done with four samples using the MinION platform, and the final sequencing of all six samples was done using the PromethION platform. Basecalling was done using Guppy v.5.0.7 and the super-accuracy model. DNA extracted using the same method was also used for short-read Illumina sequencing using the NovaSeq SP PE250 readmode.

For RNA extraction, the amoeba cells were collected by centrifugation for 5 minutes at 10000✕g at 4℃. The cells were washed with cold PAS and after another centrifugation step the cell pellets were resuspended in 1mL TRIzol Reagent (Invitrogen catalog no. 15596026). The cells were lysed by transferring the samples to Lysing Matrix E tubes (MP Biomedical cat. no. 116914500) and vortexing for 2 minutes. After transferring the homogenate to a clean tube, the samples were incubated for 5 minutes at room temperature and then centrifuged for 10 minutes at 12000✕g at 4℃ to eliminate small beads and cell debris. After transferring the supernatant to a clean tube, 200µL of Phenol/Chloroform/Isoamyl alcohol (Carl Roth cat. no. A156.2) was added, and the samples were shaken vigorously for 20 seconds. After incubation at room temperature for 2-3 minutes, the samples were centrifuged for 18 minutes at 10000✕g at 4℃. The aqueous phase was transferred to a clean tube, an equal volume of absolute ethanol (Fisher Scientific cat. no. 10644795) was added, and the samples were mixed. The samples were loaded into columns from the RNeasy Mini Kit (Qiagen cat. no. 74104) for subsequent total RNA extraction following the standard protocol. Poly(A) mRNA short read Illumina sequencing was performed using the NovaSeq SP PE150 readmode.

### Genome assembly

The long nanopore reads were basecalled using Guppy v5.0.7 (Wick et al., 2019) with the super-accuracy model, and adapter trimmed with Porechop v0.2.4 (Wick et al., 2017). The short DNA and mRNA Illumina reads were quality trimmed with FASTX-Toolkit v0.0.14 (Hannon, 2010) and PRINSEQ-lite v0.20.4 (Schmieder & Edwards, 2011). Initial long-read assemblies were done using Flye v2.9 (Kolmogorov et al., 2019), comparing the –nano-raw and –nano-hq options. These initial assemblies were compared to hybrid short and long-read assemblies using MaSuRCA v4.0.6. (Zimin et al., 2013). The overall best results were obtained using Flye v2.9 with the –nano-raw option, and these assemblies were used for downstream processing. The assemblies were manually curated by using blast-based searches against sequences of closely related organisms (Table 2), creating lists of known and unknown contigs. From the unknown lists, contaminants, contigs with a low coverage (< *Q*_1_) and short contigs (<1000 bp) were removed. The long nanopore DNA reads were mapped against the curated assemblies using Minimap2 v2.24 (H. Li, 2018, 2021) and the short Illumina DNA reads were mapped using Bowtie2 v2.5.0 (Langmead & Salzberg, 2012). Processing of the alignment files was done using SAMtools (Danecek et al., 2021). To remove spillover contaminants from the sequencing run, for each amoeba the mapping was done against the corresponding nuclear genome concatenated with all six mitochondrial genomes. The reads that mapped concordantly were extracted for separate long read re-assemblies of the nuclear and mitochondrial genomes using Flye v2.9.1, followed by sequence correction using Medaka v1.7.2 (*Medaka*, 2017/2023), followed by a polishing step using Polypolish v0.5.0 (Wick & Holt, 2022).

**Table 2.** Protein and nucleotide databases used for manual curation and gene prediction on genome assemblies.

| Species | Organism used as reference | Source and identifier |
| --- | --- | --- |
| *Acanthamoeba terricola* Neff | *Acanthamoeba terricola* Neff | RefSeq: GCF_000313135.1 |
| *Acanthamoeba castellanii* 1BU | *Acanthamoeba terricola* Neff | RefSeq: GCF_000313135.1 |
| *Acanthamoeba griffini* Sawyer | *Acanthamoeba terricola* Neff | RefSeq: GCF_000313135.1 |
| *Vermamoeba vermiformis* CDC-19 | *Vermamoeba vermiformis* | NCBI Taxonomy ID: 5778 |
| *Naegleria clarki* RU30 | *Naegleria gruberi* | RefSeq: GCF_000004985.1 |
|  | *Naegleria lovaniensis* | RefSeq: GCF_003324165.1 |
|  | *Naegleria fowleri* | RefSeq: GCF_008403515.1 |
| *Willaertia magna* T5(S)44 | *Willaertia magna* | GenBank: KX506079.1; GenBank: KX506077.1 |
|  | *Naegleria gruberi* | RefSeq: GCF_000004985.1 |
|  | *Naegleria lovaniensis* | RefSeq: GCF_003324165.1 |
|  | *Naegleria fowleri* | RefSeq: GCF_008403515.1 |

### Gene prediction and annotation

Repeats were identified and masked using RepeatModeler v2.0.4 (Flynn et al., 2020) and RepeatMaster v.4.14 (Tarailo-Graovac & Chen, 2009). The short mRNA Illumina reads were mapped against the polished genome assemblies using HISAT2 v2.2.1 (Kim et al., 2019), and Trinity v2.15.0 (Haas et al., 2013) was used to generate a genome-guided *de novo* transcriptome assembly. Transcript abundance was estimated using RSEM v1.3.3 (B. Li & Dewey, 2011), best hit isoforms were filtered with Trinity and mapped against the polished genome assemblies using GMAP v2021-12-17 (Wu & Watanabe, 2005). Processing of the alignment files was done using SAMtools (Danecek et al., 2021). Gene prediction was done using BRAKER2 (Brůna et al., 2021) using a combination of RNA-seq and protein data. For the RNA-seq data, both the full transcriptome and the filtered isoform alignments were used as input. For the protein data, we used available datasets of the same or closely related species (Table 2). Gene annotation was done using Funannotate v1.8.13 (Palmer & Stajich, 2023). The predicted proteins were fed to InterProScan v5.60 (Jones et al., 2014), eggNOG-mapper v2.1.10 (Cantalapiedra et al., 2021), Phobius v1.01 (Käll et al., 2004) and SignalP 6.0 (Teufel et al., 2022) to generate functional annotations. Ribosomal RNA and tRNA genes were annotated separately using Infernal v1.1.3 (Nawrocki & Eddy, 2013) and tRNAscan-SE v2.0.12 (Chan & Lowe, 2019), respectively. Mitochondrial genome annotation was done using the MITOS webserver (Bernt et al., 2013), followed by manual curation.

### Phylogenetic and phylogenomic analyses

Protist ribosomal RNA (rRNA) sequences with a minimum sequence length of 2000 bp were collected from the PR^2^ reference sequence database (Guillou et al., 2013). The R package *pr2database* was used to select 18S rRNA sequences from specific groups of taxa (phyla *Discosea*, *Heterolobosea* and *Tubulinea*), reference sequences of the major taxa within these groups, and sequences that are annotated in EukRibo v2 (Berney, 2022). For all amoebae sequenced, additional almost complete 18S rRNA sequences were added to cross-validate the specific strains we sequenced. Duplicate sequences were removed, and the remaining sequences were aligned with MAFFT v7.490 (Katoh & Standley, 2013) using the E-INS-i algorithm. The rRNA sequences in the nuclear genomes of the sequenced amoebae were detected using *cmsearch* within Infernal v1.1.4 (Nawrocki & Eddy, 2013) and the Rfam models RF01960, RF00002 and RF02543 for identification of 18S, 5.8S and 28S rRNA sequences, respectively. The 18S rRNA sequences were extracted from contigs that contain all rRNAs and added to the corresponding alignments using the *--addfragments* options within MAFFT. The alignments were visually checked and, if necessary, manually curated using AliView (Larsson, 2014). The alignments were filtered using Gblocks v0.91b (Castresana, 2000) (parameters -t=d, -b1=(½✕N)+1, - b2=(½✕N)+1, -b3=8, -b4=3, -b5=a, -b0=3). Phylogenetic trees of the 18S rRNA alignments were constructed using IQ-TREE v2.0.7 (Minh et al., 2020), with ModelFinder (Kalyaanamoorthy et al., 2017) to select the optimal model of sequence evolution and 1000 non-parametric bootstraps.

Phylogenomic analyses were done using PhyloFisher v1.2.11 (Tice et al., 2021), which includes a manually curated database of 240 protein-coding genes from 304 eukaryotic taxa. The standard PhyloFisher workflow was followed to collect putative homologs from the input taxa (*config.py*, *fisher.py*, *informant.py*, *working_dataset_constructor.py*) and to prepare single-protein trees (*sgt_constructor.py*). The single-protein trees were prepared for manual inspection (*forest.py*) with the standalone version of ParaSorter v1.0.4 from PhyloFisher. Orthologs and paralogs were identified during manual inspection, and orthologs were collected. The preliminary statistics after homolog collection show that the data of the newly added taxa is of good quality, which improved after manual curation of the single-protein trees (Table 3). The individual ortholog fasta files for the *Amoebozoa* and *Discoba* clades were selected (*select_taxa.py*) and processed to construct super-matrices using matrix_constructor.py within PhyloFisher. Phylogenomic trees of 228 concatenated protein gene sequences were constructed using IQ-TREE v2.2.2.7 (Minh et al., 2020), with ModelFinder (Kalyaanamoorthy et al., 2017) to select the optimal model of sequence evolution and 1000 non-parametric bootstraps.

**Table 3.**
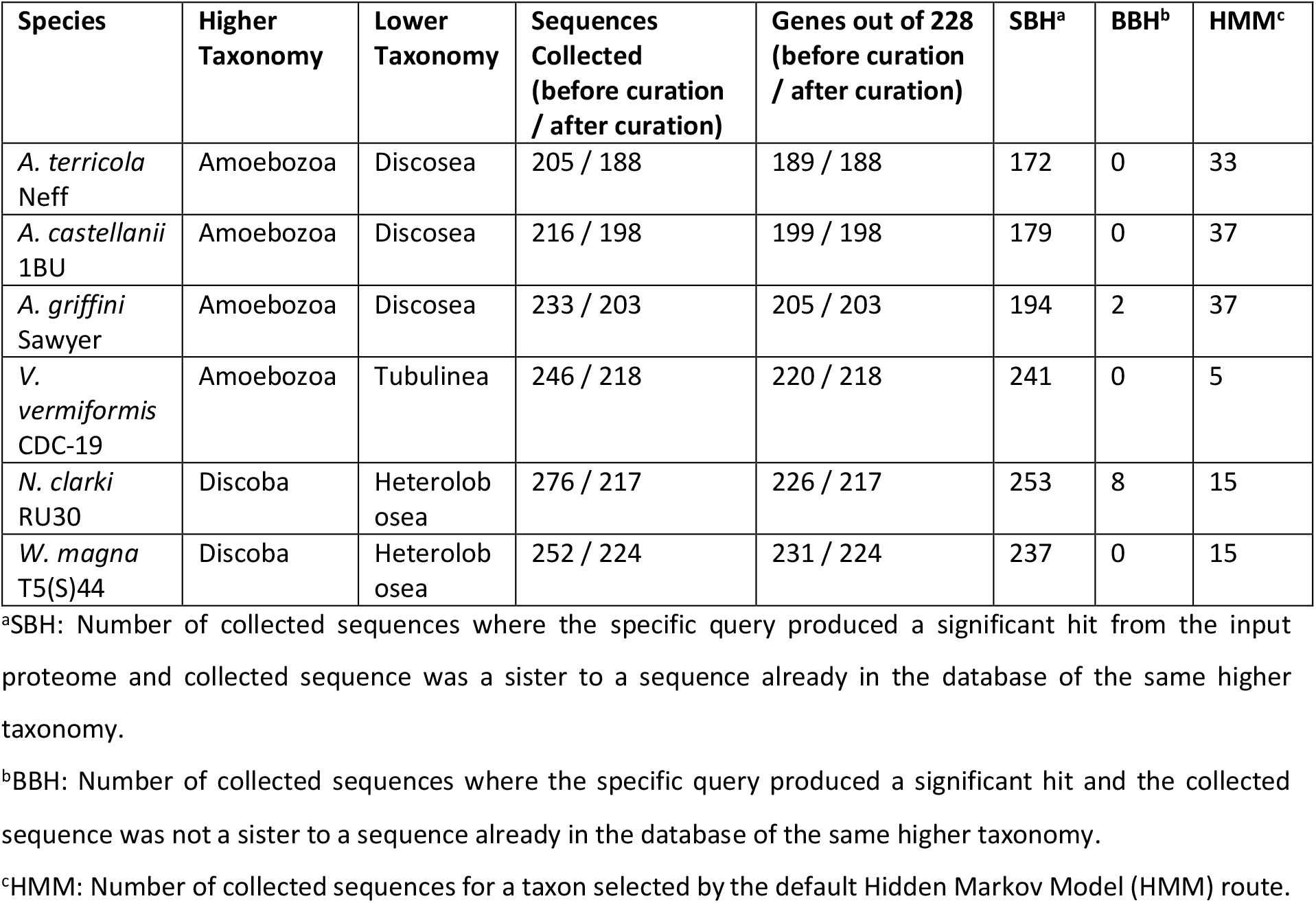
Statistics of the newly added taxa to the PhyloFisher database. Data before and after manual curation of orthologs is shown.

| Species | Higher Taxonomy | Lower Taxonomy | Sequences Collected (before curation / after curation) | Genes out of 228 (before curation / after curation) | SBH<sup>a</sup> | BBH<sup>b</sup> | HMM<sup>c</sup> |
| --- | --- | --- | --- | --- | --- | --- | --- |
| *A. terricola* Neff | Amoebozoa | Discosea | 205 / 188 | 189 / 188 | 172 | 0 | 33 |
| *A. castellanii* 1BU | Amoebozoa | Discosea | 216 / 198 | 199 / 198 | 179 | 0 | 37 |
| *A. griffini* Sawyer | Amoebozoa | Discosea | 233 / 203 | 205 / 203 | 194 | 2 | 37 |
| *V. vermiformis* CDC-19 | Amoebozoa | Tubulinea | 246 / 218 | 220 / 218 | 241 | 0 | 5 |
| *N. clarki* RU30 | Discoba | Heterolobosea | 276 / 217 | 226 / 217 | 253 | 8 | 15 |
| *W. magna* T5(S)44 | Discoba | Heterolobosea | 252 / 224 | 231 / 224 | 237 | 0 | 15 |

<sup>a</sup>SBH: Number of collected sequences where the specific query produced a significant hit from the input proteome and collected sequence was a sister to a sequence already in the database of the same higher taxonomy.
<sup>b</sup>BBH: Number of collected sequences where the specific query produced a significant hit and the collected sequence was not a sister to a sequence already in the database of the same higher taxonomy.
<sup>c</sup>HMM: Number of collected sequences for a taxon selected by the default Hidden Markov Model (HMM) route.

### Analysis of codon usage patterns

Codon usage tables, G+C composition, ENC values, CAI and COUSIN scores for coding sequences (CDSs) of the six amoebae were calculated using COUSIN v1.0 (Bourret et al., 2019). ENC quantifies codon usage in a range from extreme bias (ENC of 20: one synonymous codon is used for each amino acid) to no bias (ENC of 61: equal usage of synonymous codons) (Wright, 1990). The variation in GC content at the third codon position (GC3) accounts for much of the within-species synonymous codon usage variation in mammals (Ikemura, 1985; Aota and Ikemura, 1986), and the between-species variation in bacteria (Mute and Osawa, 1987). The relationship between GC3 and ENC values was investigated and compared to the null hypothesis (H_0_) of no translational selection. The null hypothesis was calculated as done in (Wright, 1990):

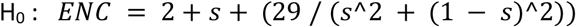

where *s* denotes GC3 scores.

The CAI score quantifies codon usage similarities between a gene and a reference set, with an index ranging from 0 to 1. If a gene always uses the most frequently used synonymous codons in the reference set, the CAI score would be 1. COUSIN also compares the codon usage preferences between a gene and a reference set but normalises the output over a null hypothesis of equal usage of synonymous codons with an index that can go below 0 or above 1. The COUSIN_18_ variant of the index considers that each of the 18 families of synonymous codons contributes equally to the global index, whereas the COUSIN_59_ variant considers that each family of synonymous codons contributes proportionally to the frequency of the corresponding amino acid in the query (Bourret et al., 2019). Giant virus genomes from the phylum *Nucleocytoviricota* (Table S4) were collected from the NCBI nucleotide database (https://www.ncbi.nlm.nih.gov/nucleotide/) in July 2021. The taxonomic classification of the collected sequences was done according to the associated GenBank information or deduced from their placement in phylogenetic trees (Koonin & Yutin, 2018; Schulz et al., 2017; Yoshikawa et al., 2019) and the International Committee on Taxonomy of Viruses (https://ictv.global/). Most sequences were full-length genomes, except for Catovirus CTV1 and Yasminevirus which are in two contigs (Table S4). Unannotated sequences were annotated using Prokka v1.14.6 (Seemann, 2014). The CDSs were extracted and used to calculate the codon usage tables and CAI and COUSIN scores at the species, genus and family levels. The Pearson product-moment correlation scores correlation between the CAI and COUSIN were calculated (Table S5). The COUSIN scores were compared using the Huber M-estimator of location and the corresponding Median Absolute Deviation (MAD) scores (Huber & Ronchetti, 2009) and Wilcoxon Rank Sum tests with continuity correction (Bauer, 1972; Hollander et al., 2015) (Table S6).

### Detection of integrated viral sequences in the amoeba genomes

To detect integrated viral sequences in the amoeba genomes, we downloaded 256 MCP genes from full-length *Nucleocytoviricota* genomes from GenBank (May 2023). We concatenated these with 196 MCP genes from virophages plus known Polinton-like viruses (PLVs) from a previous study (Bellas & Sommaruga, 2021). The total of 452 MCP genes were aligned with MAFFT v7.055b (Katoh & Standley, 2013) using the E-INS-i algorithm. HMM profiles were constructed with hmmbuild, and the amoeba genomes were interrogated with hmmsearch from HMMER v3.3.2 (http://hmmer.org/). We separately interrogated the amoeba genomes with HMM profiles from the MCPs of PLVs and virophages generated in Bellas & Sommaruga (2021), as well as with HMM profiles from five core genes (A23 packaging ATPase, D5-like helicase-primase, DNA polymerase family B, DNA/RNA helicase, poxvirus late transcription factor VLTF3-like) of *Nucleocytoviricota* generated in Schulz *et al*. (2020). The amoeba genomes were also interrogated with DIAMOND BLASTX v2.1.8.162 (Buchfink et al., 2015) using the *Nucleocytoviricota* MCP database constructed in this study and the latest MCP/PLVs/EVEs database from Bellas et al., 2023 (settings: --evalue 1e-12 --range-culling -F 15 --max-target-seqs 1, and compared with settings: --range-culling -F 15). The contigs with significant hits (Tables S7 and S8) against any of these databases were extracted using fastx_filter v.1.0 (https://github.com/amanzanom/seqTools), re-annotated with prokka v.1.14.6 (Seemann, 2014) under the kingdom viruses, and manually inspected.

### Statistics and data visualisation

The data was processed and visualised using R v.4.3.0 (R Core Team, 2023), with the packages dplyr (Wickham et al., 2023), ggplot2 (Wickham, 2016), gggenes (Wilkins, 2023), ggridges (Wilke, 2024), MASS (Venables & Ripley, 2002), viridis (Garnier et al., 2024). Phylogenetic trees were visualised using iTOL v.6.9 (Letunic & Bork, 2024). The sliding window analysis for visualising integrated viral sequences was done using bedtools v.2.30.0 (https://github.com/arq5x/bedtools2). Final figures and graphs were made with Inkscape v.1.1.2 (https://inkscape.org).

## Data availability

The raw reads (Illumina, MinION, PromethION) and the genome assemblies are available at NCBI under the following BioProject accession numbers: *Acanhtamoeba terricola* Neff: PRJNA1128888; *Acanhtamoeba castellanii* 1BU: PRJNA1128890; *Acanthamoeba griffin*i Sawyer: PRJNA1128892; *Vermamoeba vermiformis* CDC-19: PRJNA1128894; *Naegleria clarki* RU30: PRJNA1128897; and *Willaertia magna* T5(S)44: PRJNA1128895. The annotated amoeba genomes, including the curated mitochondrial genomes, nucleotide and protein alignments, distance matrices, phylogenetic and phylogenomic trees, and codon usage data of amoebae and giant viruses can be found at Zenodo https://doi.org/10.5281/zenodo.13151017.

## Acknowledgements

We would like to thank Masaharu Takemura for sharing *Acanthamoeba terricola* Neff and *Acanthamoeba castellanii medusavirus*, Julia Walochnik for sharing *Acanthamoeba castellanii* 1BU, and Bernard La Scola for sharing *Acanthamoeba griffini* Sawyer and *Willaertia magna* T5(S)44 cultures. We would also like to thank Georgi Nikolov for technical assistance. The computational results of this work have been achieved using the Life Science Compute Cluster (LiSC) of the University of Vienna. This project has received funding from the European Union’s Horizon 2020 research and innovation programme under the Marie Sklodowska-Curie grant agreement No. 891572 and the European Union (ERC, CHIMERA, 101039843). Views and opinions expressed are however those of the author(s) only and do not necessarily reflect those of the European Union or the European Research Council Executive Agency. Neither the European Union nor the granting authority can be held responsible for them. AW and MH acknowledge funding from the Austrian Science Fund Cluster of Excellence “microplanet” [doi.org/10.55776/COE7].

