## Supplementary material for "Novel high-quality amoeba genomes reveal widespread codon usage mismatch between giant viruses and their hosts": Figures S1-10

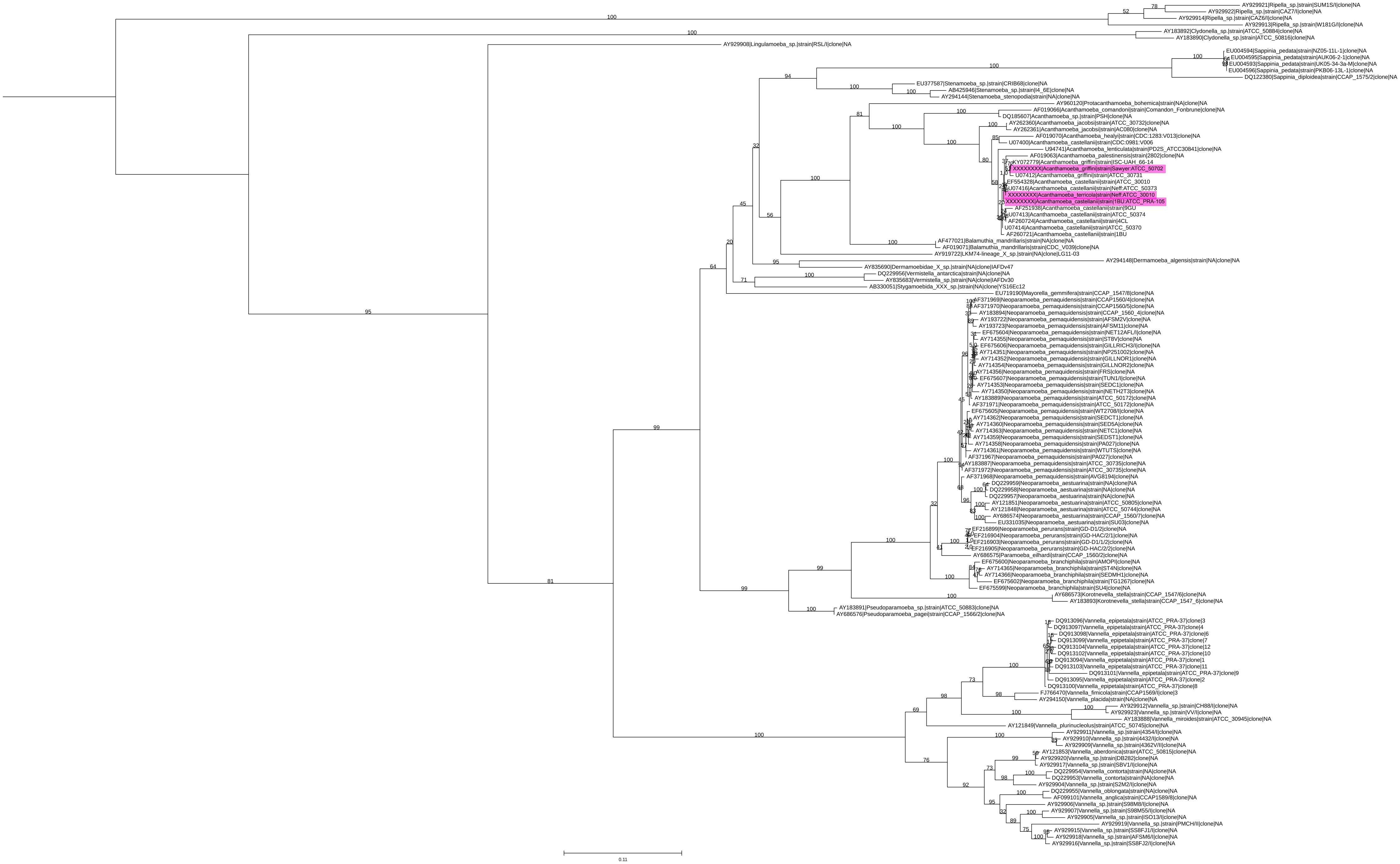


**Figure S1**. Phylogenetic tree of the 18S rRNA sequences of the phylum *Discosea*. This maximum likelihood tree was constructed on a filtered nucleotide alignment under the GTR+F+R5 model of evolution with counted frequencies and 1000 non-parametric boopstrap replicates. The tree is rooted using midpoint rooting. The added taxa are highlighted in pink.

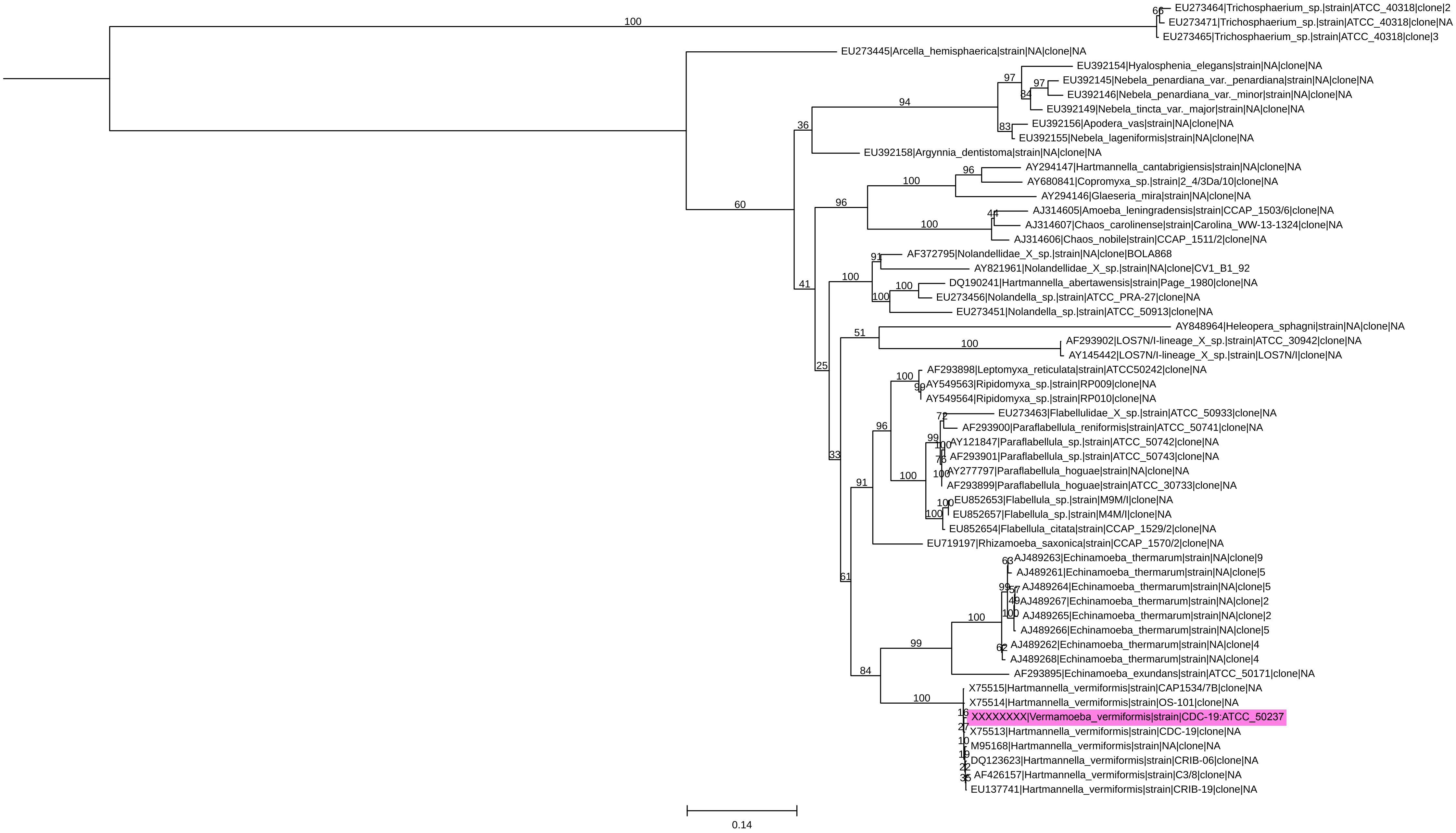


**Figure S2**. Phylogenetic tree of the 18S rRNA sequences of the phylum *Tubulinea*. This maximum likelihood tree was constructed on a filtered nucleotide alignment under the TIM2+F+I+G4 model of evolution with counted frequencies and 1000 non-parametric boopstrap replicates. The tree is rooted using midpoint rooting. The added taxon is highlighted in pink.

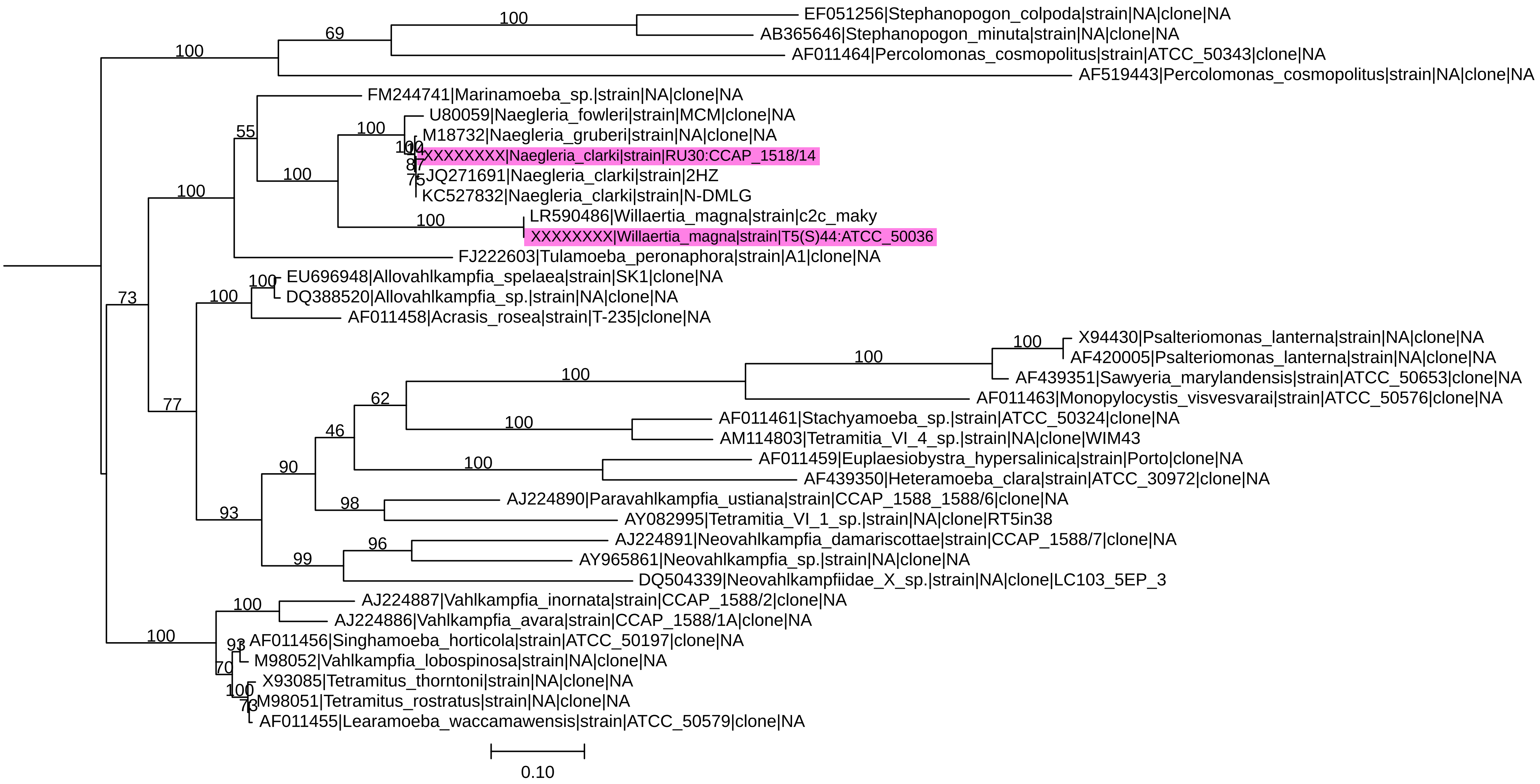


**Figure S3**. Phylogenetic tree of the 18S rRNA sequences of the phylum *Heterolobosea*. This maximum likelihood tree was constructed on a filtered nucleotide alignment under the TIM2+F+R4 model of evolution with counted frequencies and 1000 non-parametric boopstrap replicates. The tree is rooted using midpoint rooting. The added taxon is highlighted in pink.

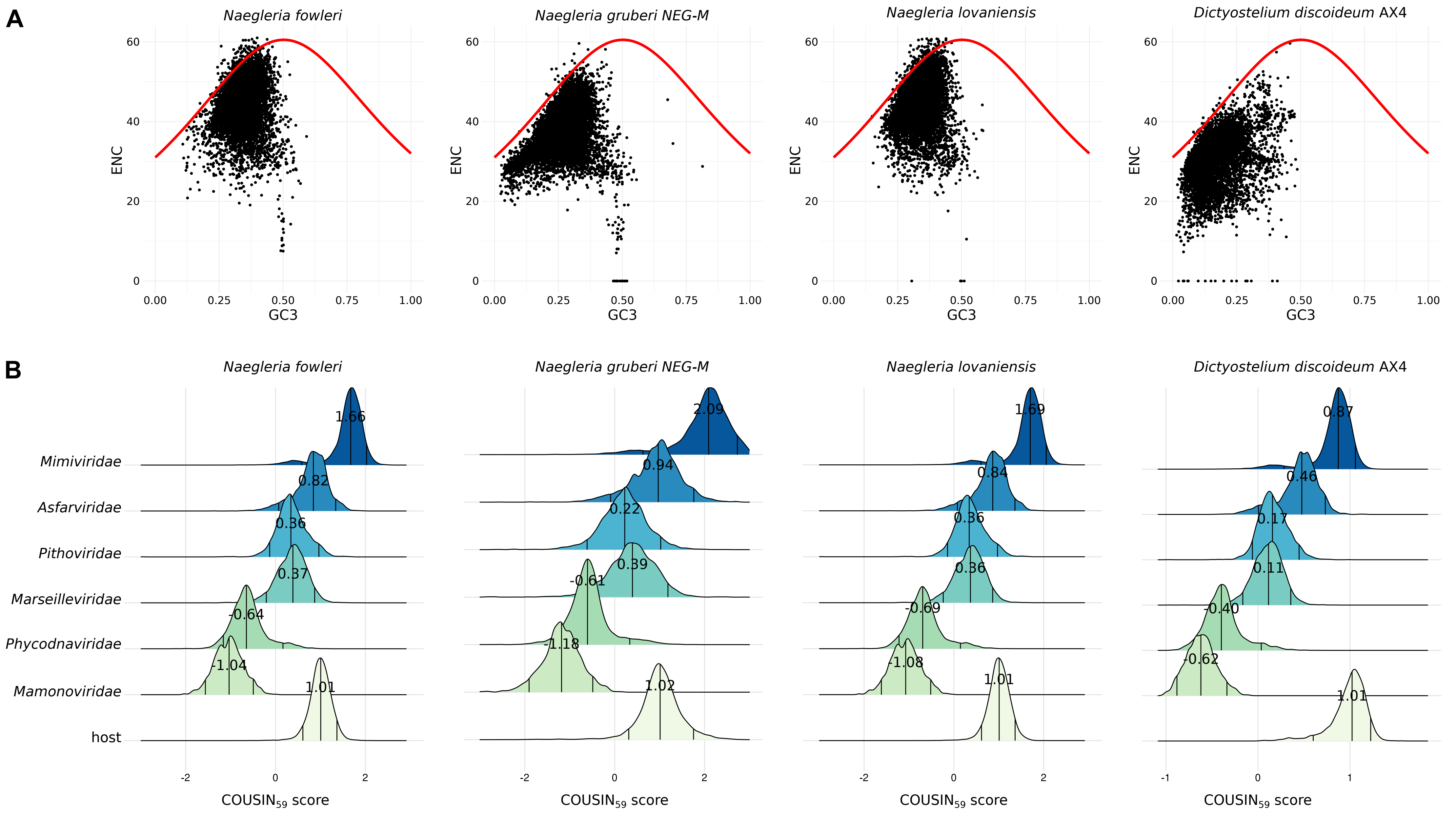


**Figure S4**. Codon usage preferences of giant viruses with respect to previously published high quality amoebae genomes (**A**) Codon usage patterns for amoebae from the *Discoba* (*N. fowleri*, *N. gruberi*, *N. lovaniensis*) and *Amoebozoa* (*D. discoideum*) eukaryotic clades. The ENC values were plotted against GC content at the third codon position (GC3). Each black dot represents a gene. The continuous red curves represent the relationships between ENC and GC3 under the null hypothesis of no translational selection. If a particular gene lies on or just below the red curve, it is suggested to be subjected to mutational bias only (i.e. G+C compositional constraints). (**B**) Density curves of the COUSIN59 score for giant viruses relative to their known and possible hosts. The density curves are separated by viral family (*Mimiviridae*: $N_{viruses}$=29, $N_{CDS}$=28969, *Asfarviridae*: $N_{viruses}$=9, $N_{CDS}$=4292, *Pithoviridae*: $N_{viruses}$=6, $N_{CDS}$=3393, *Marseilleviridae*: $N_{viruses}$=10, $N_{CDS}$=5735, *Phycodnaviridae*: $N_{viruses}$=11, $N_{CDS}$=9236, *Mamonoviridae*: $N_{viruses}$=2, $N_{CDS}$=890). The bottom density curve always indicates the scores for the host for which the name is indicated on top of each plot. The three lines within the density curves indicate the 95% confidence interval. Viruses that have their density curves within the host distribution, have similar codon usage preferences to their hosts. The numbers within each curve indicate the center values as estimated by the Huber M-estimator of location. The COUSIN scores can be interpreted as follows: a score of 1 indicates that the codon usage preferences of viruses are similar to those of the corresponding host; a score of 0 indicates that there is equal usage of synonymous codons; above 1 indicates that codon usage preferences are similar but of larger magnitude (meaning that the codons that are most frequently used in the host are used even more frequently in the virus); between 0 and 1 indicates codon usage preferences are similar but of smaller magnitude (meaning that the codons that are less frequently used in the host are used even less frequently in the virus); below 0 means that the codon usage preferences of viruses are opposite to those of the corresponding host.

### *Acanthamoeba terricola* Neff

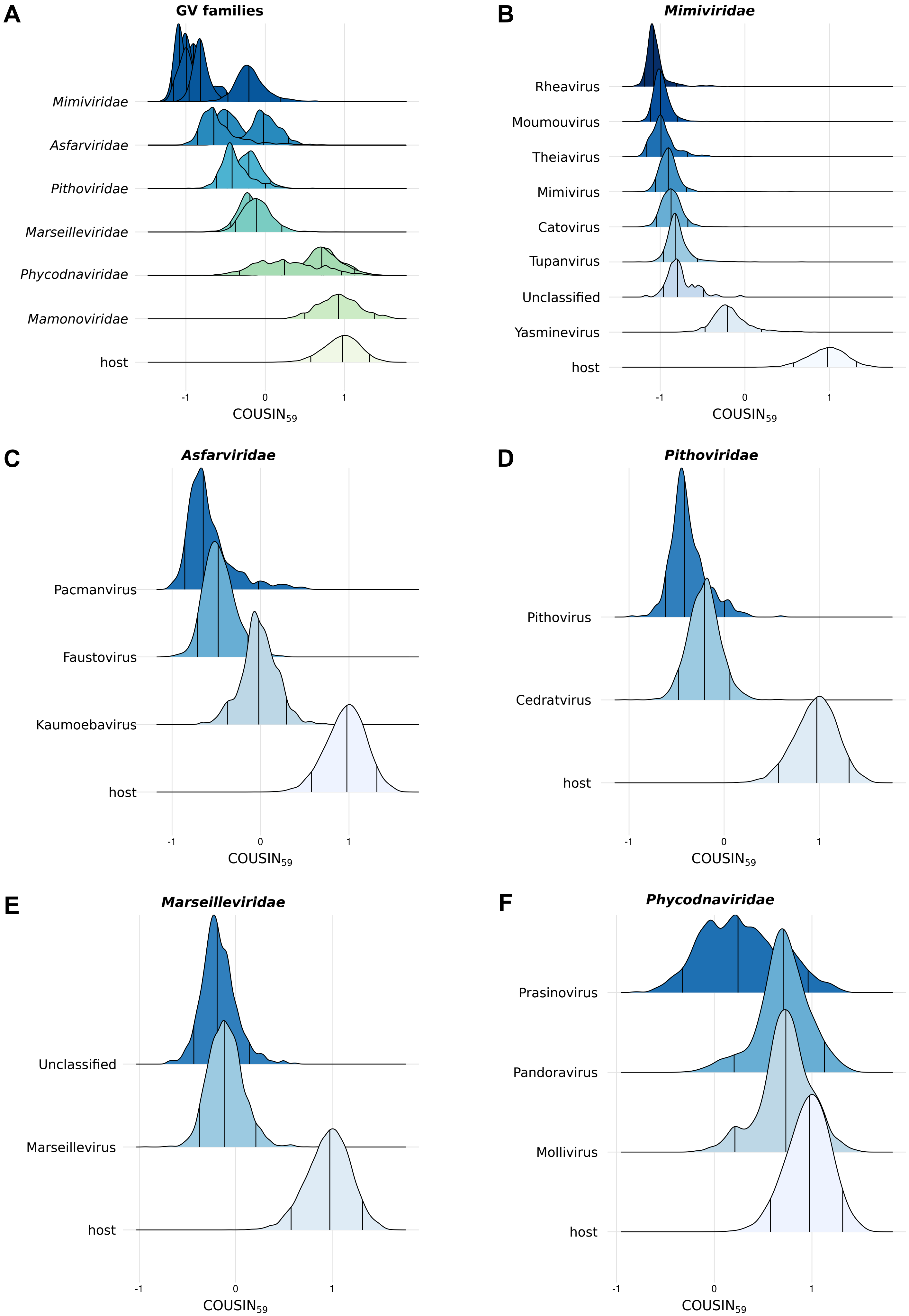


**Figure S5**. Density curves of the $COUSIN_{59}$ score for giant viruses relative to the host *Acanthamoeba terricola* Neff. (**A**) An overview of all viral families included (y-axis). The density curves are separated by viral genera. (**B-F**) Each pot represents a zoom of one viral family where the genera are separated on the y-axis. The bottom density curve always indicates the scores for the host. The three lines within the density curves indicate the 95% confidence interval. Viruses that have their density curves within the host distribution, have similar codon usage preferences to their hosts.

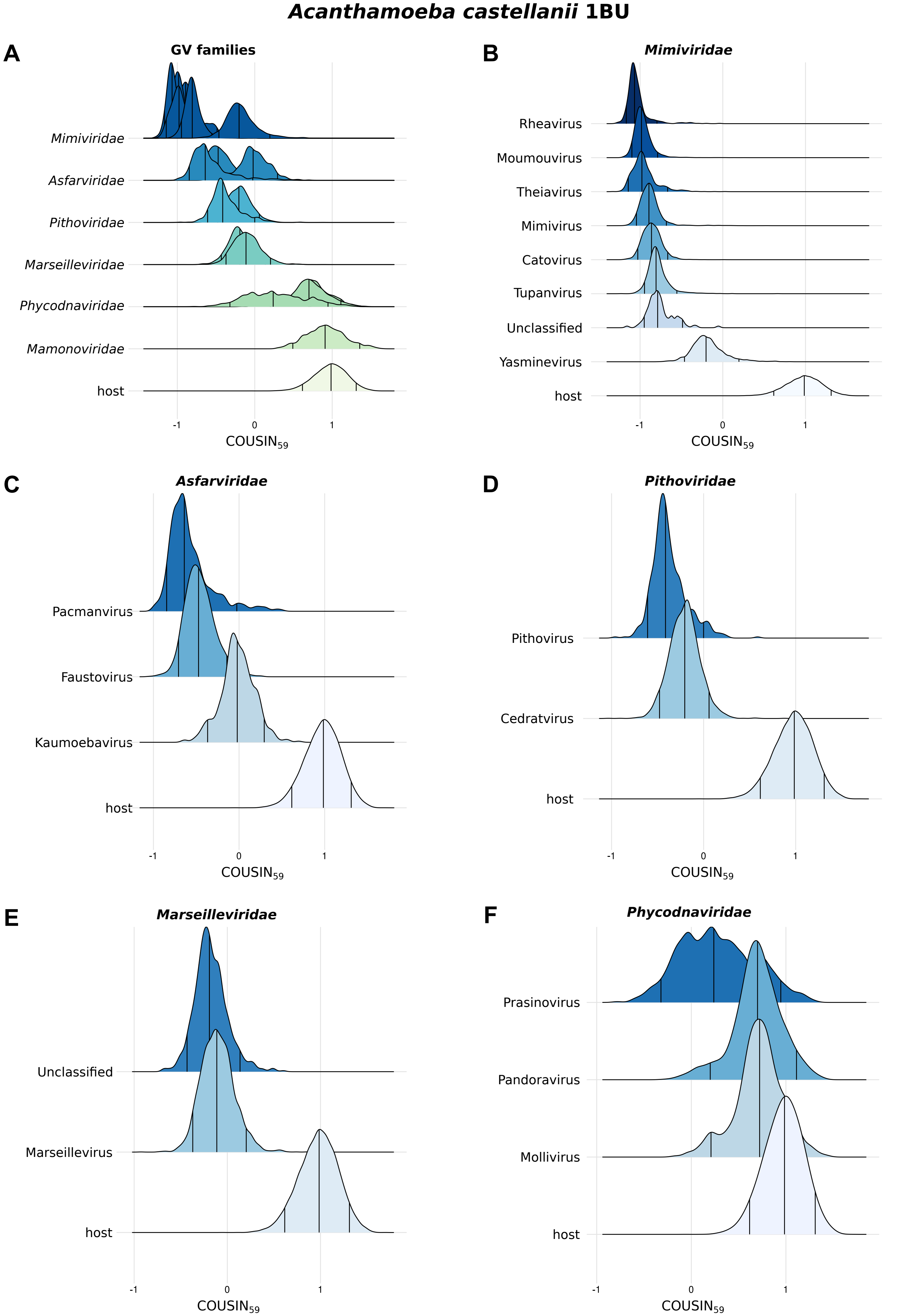


**Figure S6**. Density curves of the $COUSIN_{59}$ score for giant viruses relative to the host *Acanthamoeba castellanii* 1BU. (**A**) An overview of all viral families included (y-axis). The density curves are separated by viral genera. (**B-F**) Each pot represents a zoom of one viral family where the genera are separated on the y-axis. The bottom density curve always indicates the scores for the host. The three lines within the density curves indicate the 95% confidence interval. Viruses that have their density curves within the host distribution, have similar codon usage preferences to their hosts.

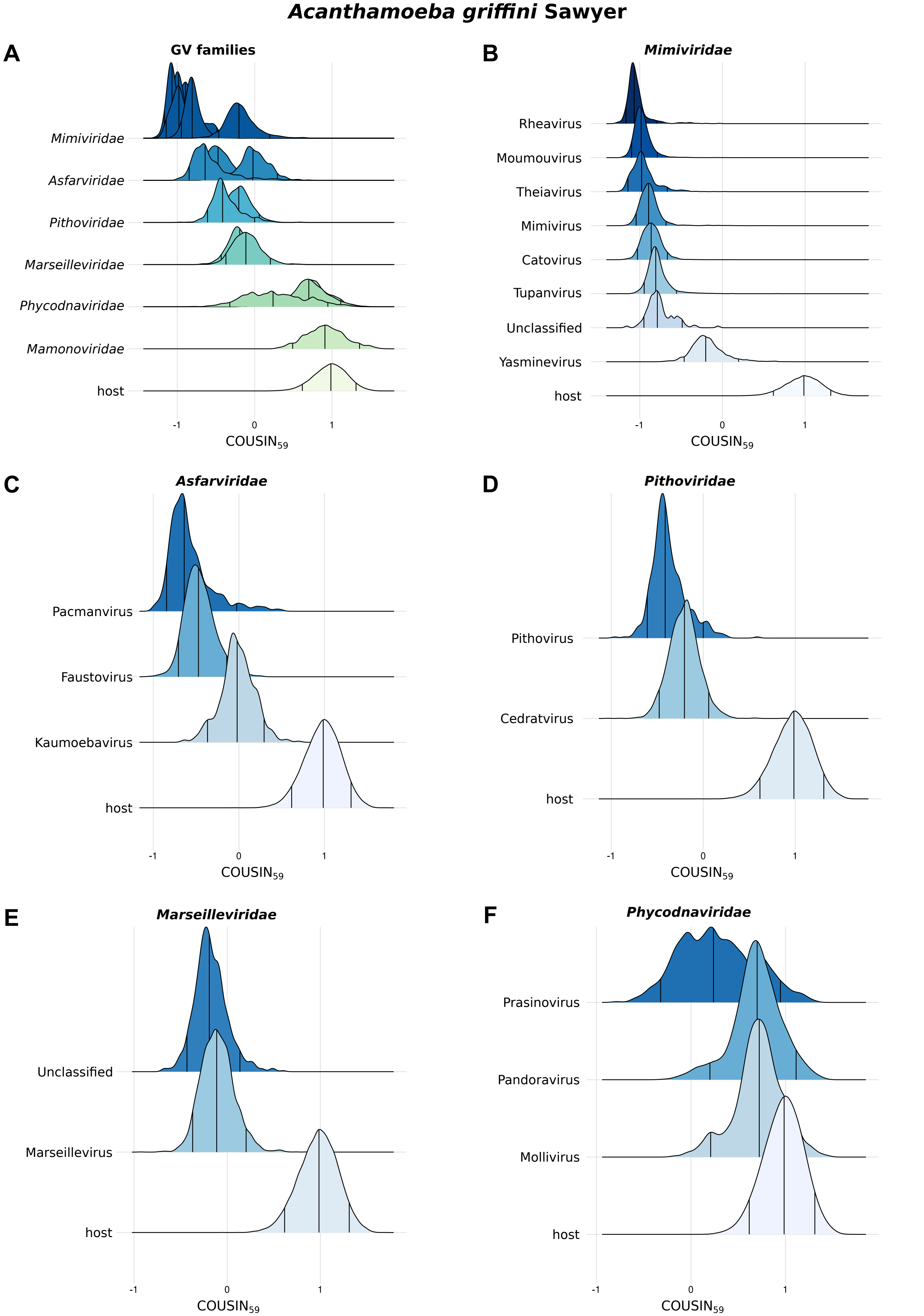


**Figure S7**. Density curves of the $COUSIN_{59}$ score for giant viruses relative to the host *Acanthamoeba griffini* Sawyer. (**A**) An overview of all viral families included (y-axis). The density curves are separated by viral genera. (**B-F**) Each pot represents a zoom of one viral family where the genera are separated on the y-axis. The bottom density curve always indicates the scores for the host. The three lines within the density curves indicate the 95% confidence interval. Viruses that have their density curves within the host distribution, have similar codon usage preferences to their hosts.

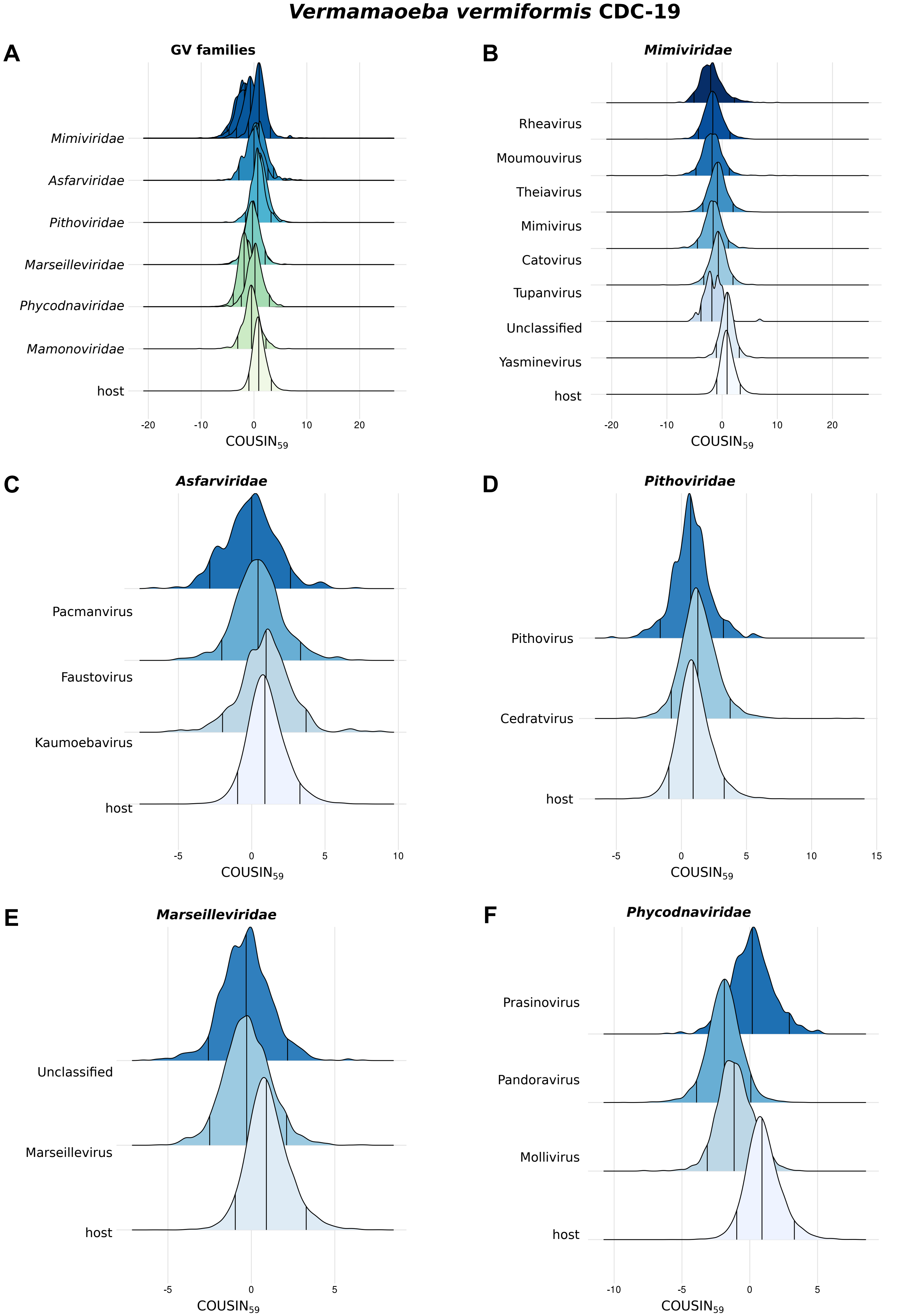


**Figure S8**. Density curves of the $COUSIN_{59}$ score for giant viruses relative to the host *Vermamoeba vermiformis* CDC-19. (**A**) An overview of all viral families included (y-axis). The density curves are separated by viral genera. (**B-F**) Each pot represents a zoom of one viral family where the genera are separated on the y-axis. The bottom density curve always indicates the scores for the host. The three lines within the density curves indicate the 95% confidence interval. Viruses that have their density curves within the host distribution, have similar codon usage preferences to their hosts.

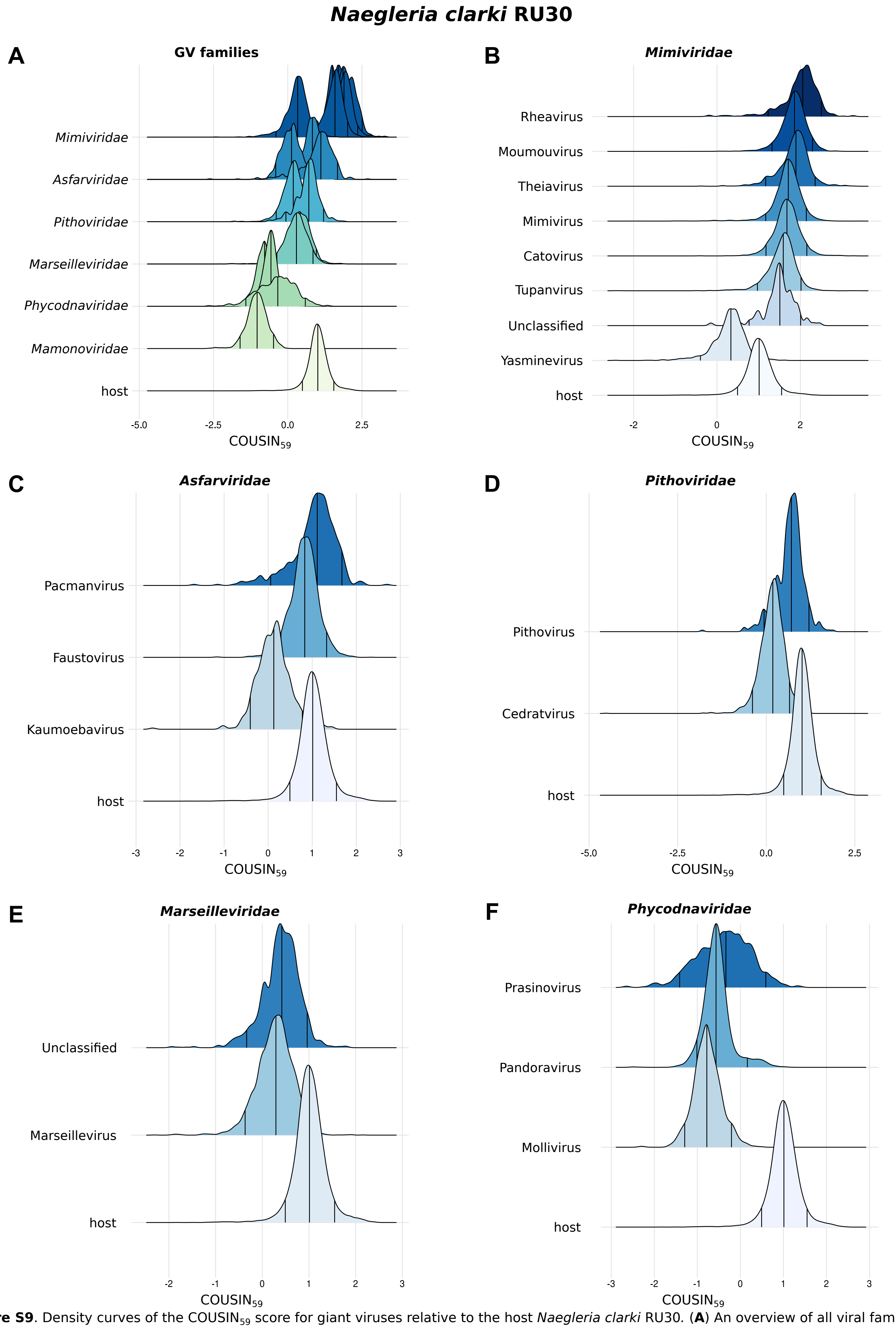


**Figure S9**. Density curves of the $COUSIN_{59}$ score for giant viruses relative to the host *Naegleria clarki* RU30. (**A**) An overview of all viral families included (y-axis). The density curves are separated by viral genera. (**B-F**) Each pot represents a zoom of one viral family where the genera are separated on the y-axis. The bottom density curve always indicates the scores for the host. The three lines within the density curves indicate the 95% confidence interval. Viruses that have their density curves within the host distribution, have similar codon usage preferences to their hosts.

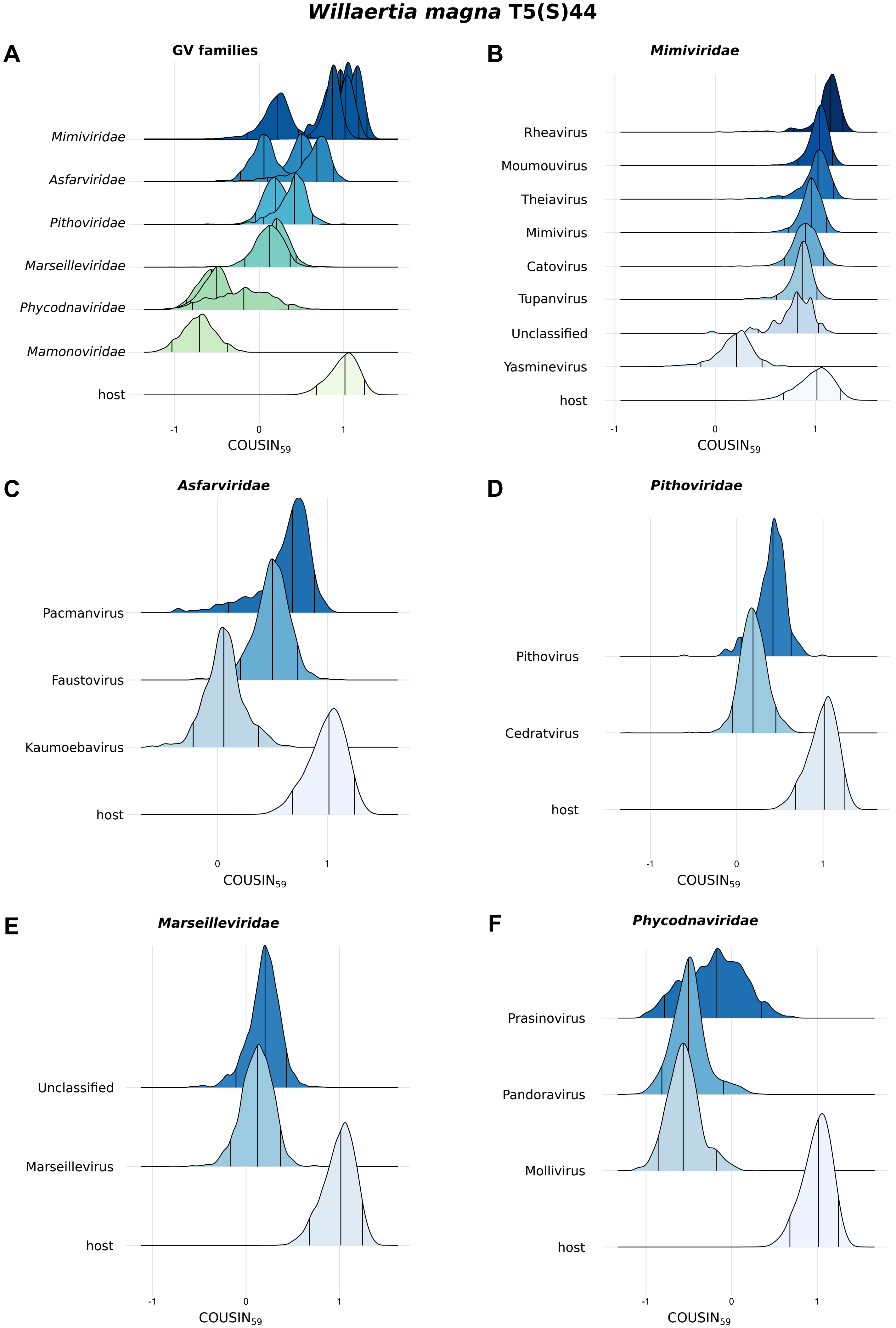


**Figure S10**. Density curves of the $COUSIN_{59}$ score for giant viruses relative to the host *Willaertia magna* T5(S)44. (**A**) An overview of all viral families included (y-axis). The density curves are separated by viral genera. (**B-F**) Each pot represents a zoom of one viral family where the genera are separated on the y-axis. The bottom density curve always indicates the scores for the host. The three lines within the density curves indicate the 95% confidence interval. Viruses that have their density curves within the host distribution, have similar codon usage preferences to their hosts.
